# Enteric glutamatergic interneurons regulate intestinal motility

**DOI:** 10.1101/2024.03.24.586153

**Authors:** Ryan Hamnett, Jacqueline L. Bendrick, Keiramarie Robertson, Eric Tianjiao Zhao, Julia A. Kaltschmidt

## Abstract

The enteric nervous system (ENS) controls digestion autonomously via a complex neural network within the gut wall. Enteric neurons expressing glutamate have been identified by transcriptomic studies as a distinct subpopulation, and glutamate can affect intestinal motility by modulating enteric neuron activity. However, the nature of glutamatergic neurons, their position within the ENS circuit, and their function in regulating gut motility are unknown.

Here, we identify glutamatergic neurons as longitudinally projecting descending interneurons in the small intestine and colon, in addition to a novel class of circumferential neurons only in the colon. Both populations make synaptic contact with diverse neuronal subtypes, and signal with a variety of neurotransmitters and neuropeptides in addition to glutamate, including acetylcholine and enkephalin. Knocking out the glutamate transporter VGLUT2 from enkephalin neurons profoundly disrupts gastrointestinal transit, while *ex vivo* optogenetic stimulation of glutamatergic neurons initiates propulsive motility in the colon. This motility effect is reproduced when stimulating only the descending interneuron class, marked by Calb1 expression. Our results posit glutamatergic neurons as key interneurons that regulate intestinal motility.

## Introduction

Digestion is regulated by the enteric nervous system (ENS), an intrinsic network of neuronal circuits within the gut wall that acts independent of the central nervous system (CNS)^1^. Intestinal motility is typically initiated by sensory neurons detecting luminal contents and stimulating motor neuron activity via interneurons^2–4^. Each neuron type is further divided into subpopulations that have specific roles and express unique combinations of neurotransmitters and neuropeptides, enabling the full spectrum of gastrointestinal (GI) functions^5,6^.

Glutamate is expressed in enteric neurons, typically alongside the more widespread acetylcholine (ACh), and can mediate excitatory postsynaptic potentials in enteric neurons via myriad ionotropic and metabotropic receptors to influence GI motility^7–11^. Furthermore, glutamate may be involved in several pathologies of the GI tract, including inflammatory bowel disease (IBD) and ischemia/reperfusion injury^12–15^. Recent single cell (sc)RNAseq studies identified glutamate in only 2-3 enteric neuron subpopulations based on VGLUT2 expression, the predominant glutamatergic marker in the ENS^5,6,16,17^. However, the expression profiles of these clusters do not match the typical neurochemical coding of known functional enteric subtypes^5,6,17,18^. How glutamate achieves its effects and through which recipient neurons are also little explored, though previous studies show that two morphologically-distinct ENS subtypes, Dogiel Type 1 and Type 2 neurons, respond to exogenous glutamate, which, combined with widespread expression of glutamate receptors, suggests glutamatergic neurons communicate with large portions of the enteric circuit^9,12,19^.

Here, we use intersectional genetics, virally mediated single neuron tracing, immunohistochemistry (IHC) and optogenetics to elucidate the role of glutamatergic neurons in GI motility and explore their position within the enteric circuit. In the small intestine (SI) and colon, glutamatergic neurons are descending interneurons that project over several centimetres, while an additional morphological class exists in the colon, which project circumferentially. Both populations form putative synapses with numerous neuronal subpopulations, including calretinin, secretagogin and somatostatin neurons. Knocking out VGLUT2 from most enteric glutamatergic neurons profoundly quickens total GI transit *in vivo*, while optogenetically activating these neurons *ex vivo* initiates and accelerates propulsive motility of faecal contents in the colon. This propulsive effect is reproduced when exciting only Calb1 neurons, which represent the longitudinal descending interneuron glutamatergic subpopulation. Together, our results suggest at least two morphologically distinct glutamatergic populations, capable of engaging multiple diverse components of the enteric circuit to stimulate GI motility.

## Results

### Neurochemical coding of VGLUT2 neurons suggests excitatory, non-motor identity

VGLUT2 is expressed at varying levels in all regions of the mouse myenteric plexus (MP): the duodenum, jejunum, ileum, and proximal and distal colon^20^. We sought to validate and extend recent scRNAseq findings of VGLUT2 co-expression with other ENS markers using immunohistochemistry (IHC) and RNAscope to provide insight into the neurochemical coding and therefore potential function of VGLUT2 neurons. Given that VGLUT2 protein itself is not typically found in the soma^11^, making co-expression with other markers difficult to ascertain, we used VGLUT2 recombinase lines (VGLUT2-Cre or VGLUT2-Flp) crossed with reporters (Cre-dependent tdTomato or Flp-dependent GFP) to visualise VGLUT2 neurons as VGLUT2^tdT^ and VGLUT2^GFP^, respectively.

We observed VGLUT2 somata only in the MP, though some VGLUT2^GFP^ fibres were seen in the submucosal plexus (SMP; Extended Data Fig. 1a). In the MP, almost all VGLUT2 neurons were cholinergic, ∼30-50% were serotonergic, depending on region, and almost no VGLUT2 neurons co-expressed nNOS (Fig. 1a-c,i). Despite this cholinergic identity, only 10-16% of VGLUT2 neurons co-expressed calretinin, another marker that is predominantly cholinergic, and which delineates ascending interneurons and excitatory motor neurons (Fig. 1d,j). Secretagogin, a calcium binding protein currently of unknown function in the ENS, colocalised with VGLUT2 significantly more strongly in the SI compared to the colon (Fig. 1e,j), suggesting a difference in the types of VGLUT2 neurons found between the two organs. This was similarly observed for co-expression between Vip^tdT^ and VGLUT2^GFP^, which was significantly less in the colon (Fig. 1f,k). VIP is known to mark inhibitory motor neurons together with nNOS, as well as descending interneurons. Given that VGLUT2 does not colocalise with nNOS, it is likely that VGLUT2+/VIP+ cells are interneurons. Penk^tdT^ co-expressed with more than 50% of VGLUT2^GFP^ cells across most regions of the intestines (Fig. 1g,k). Penk^tdT^ is a marker of enkephalin neurons, which are typically interneurons or excitatory motor neurons. Finally, *Calcb*, the mRNA of CGRP, which has previously been used as a sensory neuron marker, colocalised with 10-20% of VGLUT2 neurons in the ileum and colon, but not at all in the proximal SI (Fig. 1h,k).

**Fig. 1.**
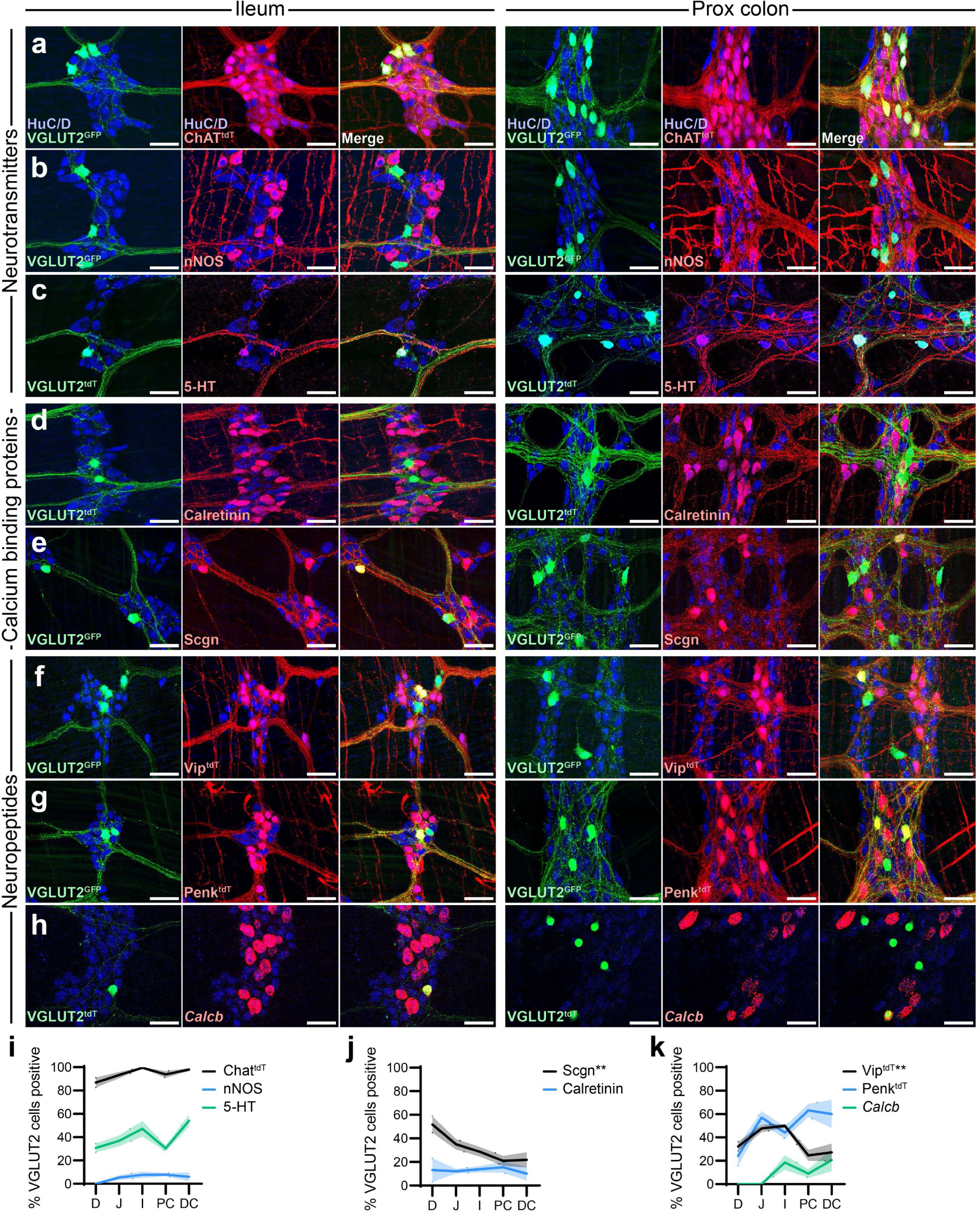
Glutamatergic neuron coexpression with other ENS markers. **a-h**, Representative images of adult wholemount MP showing VGLUT2^tdT^ or VGLUT2^GFP^ (green) and neuronal label HuC/D (blue) alongside immunohistochemical labels or genetically encoded reporters for neurotransmitters (red, **a-c**), calcium binding proteins (red, **d,e**), and neuropeptides (red, **f-h**) in the ileum (left) and proximal colon (right). Scale bars: 50 µm. **i-k**, Proportion of VGLUT2^tdT^ or VGLUT2^GFP^ neurons (mean ± SEM) positive for each neuronal marker across intestinal regions as in **a-h**, divided into neurotransmitters (**i**), calcium binding proteins (**j**) and neuropeptides (**k**). n=3-9. All tests one-way ANOVA to determine differences for a single marker colocalising with VGLUT2^tdT^ or VGLUT2^GFP^ across intestinal regions. **p< 0.01.

These data suggest that VGLUT2 is likely to be expressed in interneurons, given its coexpression with ChAT, 5-HT, VIP and Penk, while it is unlikely to be in motor neurons due to its lack of colocalisation with nNOS and calretinin, and the absence of any fibres in the smooth muscle layers.

### VGLUT2 neurons display two different morphologies in the colon

To further establish VGLUT2 neuron identity beyond their neurochemical coding, we employed a sparse labelling strategy to visualise the morphology of individual VGLUT2 neurons^21^. By injecting low titre Cre-dependent AAV-GFP systemically into the retro-orbital sinus of VGLUT2-Cre mice (VGLUT2^AAV-GFP^), a small number of enteric neurons are transduced and will express GFP, allowing visualisation of their full morphology without the fibres of other neurons interfering.

We observed two distinct morphologies of VGLUT2^AAV-GFP^ neurons: longitudinal descending interneurons in both the SI and colon (Fig. 2a,c,d; VGLUT2^Long^), and circumferential neurons (VGLUT2^Circ^), which were found only in the colon (Fig. 2b,e) and to our knowledge represent a previously undescribed morphology. VGLUT2^Long^ neurons always projected aborally over distances of up to 40 mm (Fig. 2f,g), were monoaxonal, and had sparse branches along the length of the primary fibre, though the primary fibre sometimes bifurcated, particularly in the colon (Fig. 2a,c,d). These longitudinal neurons tended to be longer in the SI than colon (Fig. 2h). In contrast, VGLUT2^Circ^ neurons projected in either direction around the circumference of the colon, largely staying within a one or two ganglionic stripes (Fig. 2b,f)^20^. VGLUT2^Circ^ neurons often projected over the anti-mesenteric border, though never over the mesenteric border, and they arborised in myenteric ganglia, which we term ‘nests’ (Fig. 2b,e). Circumferential neurons also had one main axon, though they had several small, short filaments near the soma (Fig. 2b, inset), distinguishing them from previously described circumferential neurons^22^. VGLUT2^Circ^ were significantly shorter than VGLUT2^Long^, and significantly more branched (Fig. 2h,i).

**Fig. 2.**
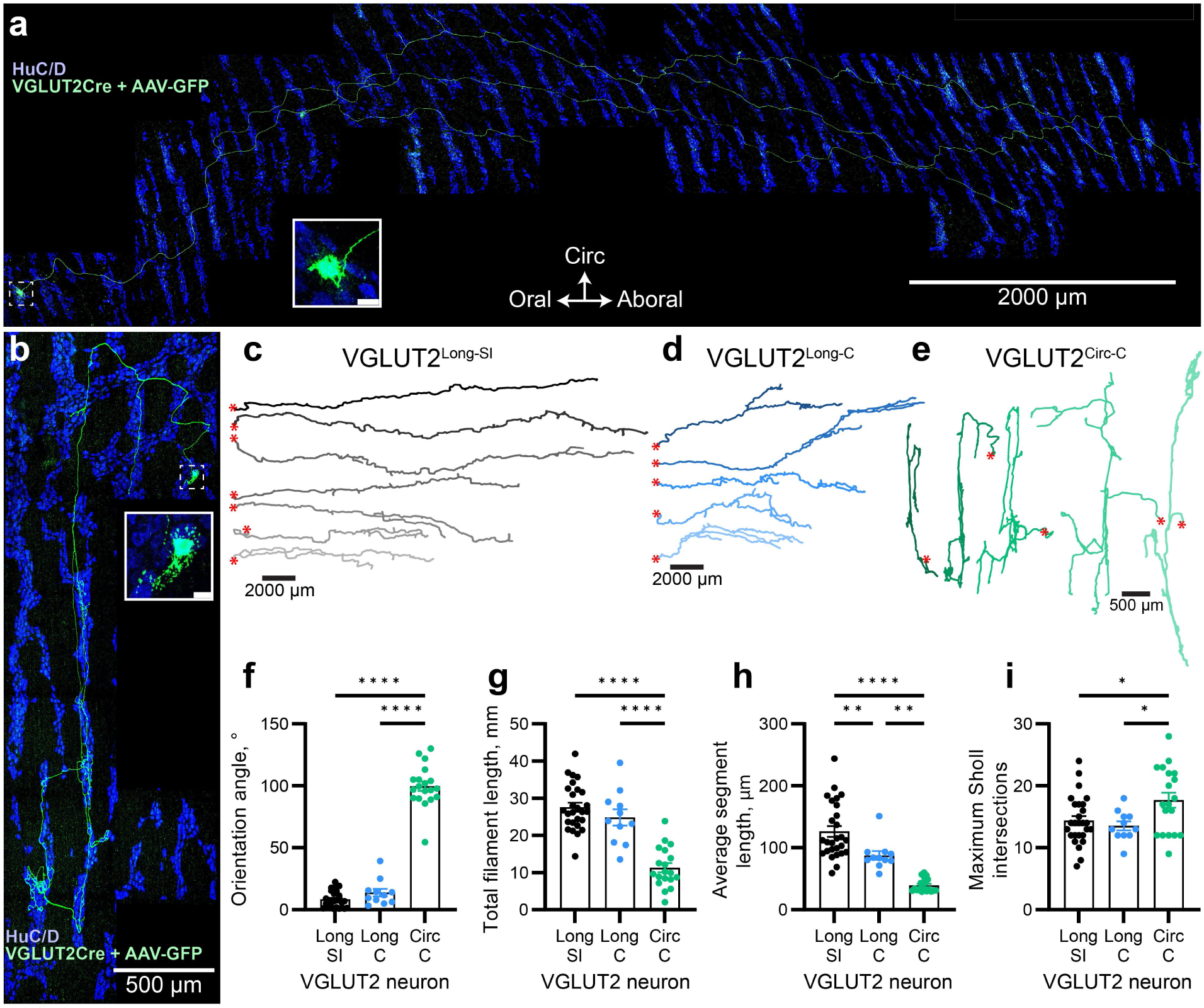
Glutamatergic neurons are divided into 2 morphological classes. **a,b**, Representative images of VGLUT2-Cre neurons transduced by Cre-dependent AAV-GFP and immunostained for GFP (green) to reveal full neuron morphology, alongside HuC/D (blue). Longitudinal (**a**) and circumferential (**b**) neurons are shown. Neuronal somata are inset (white boxes). Scale bars as indicated. **c-e**, Representative traces of AAV-GFP-labelled longitudinal neurons in the small intestine (**c**) and colon (**d**), and circumferential neurons in the colon (**e**). Soma location indicated by red asterisk. **f-i**, Quantification (mean ± SEM) of VGLUT2^AAV-GFP^ neuron orientation (**f**), total filament length (**g**), average segment length (**h**), and maximum Sholl intersections (**i**). n=11-27. Each dot represents a different neuron, taken from across 7 mice. Abbreviations: C: colon; SI: small intestine.

Knowing that VGLUT2 broadly colocalises with a number of ENS markers from IHC (Fig. 1) and from scRNAseq studies^5,6^, we next asked in which morphological class of VGLUT2 neurons (VGLUT2^Long^ or VGLUT2^Circ^) the expression of a given marker occurs, by performing the same viral tracing approach with other Cre reporter lines. We additionally asked if these markers were only expressed in VGLUT2-like descending longitudinal and circumferential neurons, or if they also represented other enteric neuron classifications.

We chose markers that were expected to label diverse populations and that had available genetic tools: Penk^AAV-GFP^, Tac1^AAV-GFP^ and Vip^AAV-GFP^ (Extended Data Fig. 1a-c). The neurons were assigned functional classifications based on morphological measurements such as orientation, length, branching, and fibre location (e.g. within muscle for motor neurons)(Extended Data Fig. 1d-i). Penk^AAV-GFP^ neurons were classified as ascending interneurons, descending interneurons (including some that bore different orientation and branching patterns to VGLUT2^Long^), excitatory motor neurons, and circumferential neurons (Extended Data Fig. 2j,m). Tac1^AAV-GFP^ showed morphology consistent with ascending interneurons, excitatory motor neurons, and circumferential neurons (Extended Data Fig. 2k,n). Finally, Vip^AAV-GFP^ neurons were predominantly inhibitory motor neurons in the SI with some descending interneurons, while the colon additionally had ascending interneurons and neurons that appeared to project to the epithelium (Extended Data Fig. 2l,o). The circumferential neurons seen in both Penk^AAV-GFP^ and Tac1^AAV-GFP^ resembled VGLUT2^Circ^ morphology, while Vip^AAV-GFP^ and some Penk^AAV-GFP^ descending interneurons resembled VGLUT2^Long^.

Thus, sparse viral labelling revealed the projection patterns of VGLUT2 longitudinal and circumferential neurons, as well as Penk, Tac1 and Vip neurons, a proportion of which shared the same morphology as VGLUT2 neurons. 3 to 6 different neuronal classifications were observed for Penk, Tac1 and Vip, compared to 2 for VGLUT2, suggesting the relative specificity of VGLUT2 as a marker for enteric neuron subtypes.

### VGLUT2 neurons form synaptic varicosities along entire length of fibre

To visualise where individual VGLUT2 neurons form synapses, we expressed synaptophysin-fused tdTomato in VGLUT2-Cre neurons (VGLUT2^syn-tdT^) to highlight putative pre-synaptic varicosities. We combined this genetic strategy with sparse viral labelling, using Cre-dependent AAV-GFP as previously described, to enable us to assign varicosities to specific regions of neurons of known morphology.

Given the long length yet sparse branching of VGLUT2^Long^ neurons, we hypothesised that they would form synapses along the entire length of the fibre, not just on the branches. Confirming this, GFP+ varicosities colocalised with syn-tdT on both the primary fibre and the branches of VGLUT2^Long^ neurons (Fig. 3a-c). For VGLUT2^Circ^ neurons, varicosities were found within nests in myenteric ganglia (Fig. 3d). Varicosities were far denser on branches and nests than they were on primary fibres (Fig. 3e,i), and more likely to be glutamatergic, evidenced by colocalisation with VGLUT2 IHC (Fig. 3f,j), with the exception of the terminal region of the fibre of VGLUT2^Long^ neurons (Fig. 3b). Glutamatergic varicosities on branches were also far more likely to be in close contact with other HuC/D+ neurons (Fig. 3g,k), ∼10% of which were other VGLUT2 neurons (Fig. 3h,l). Despite the high proportion of glutamatergic synapses at the primary fibre terminal, only ∼50% appeared to contact a HuC/D+ cell (Fig. 3g,k).

**Fig. 3.**
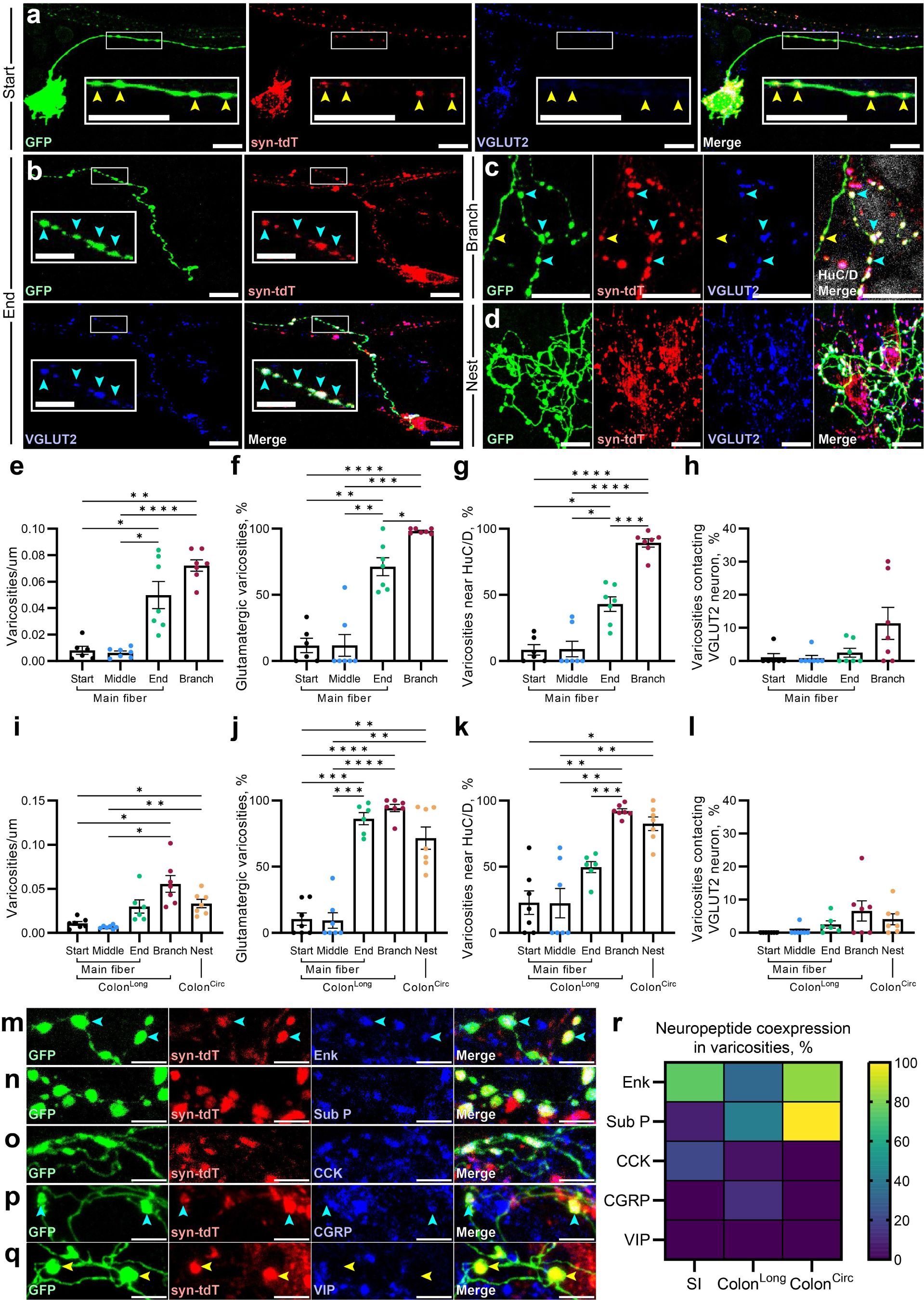
Putative synaptic varicosities are found in all regions of VGLUT2 neurons. **a-d**, Representative images of VGLUT2^syn-tdT^ (red) neurons transduced by Cre-dependent AAV-GFP (green) and immunolabelled for VGLUT2 IHC (blue) to identify putative synapses at the start (**a**), end (**b**) and on the branches (**c**) of VGLUT2^Long^ neurons, and in nests in VGLUT2^Circ^ neurons (**d**). Inset white boxes indicate zoomed areas (**a,b**). Yellow arrows indicate GFP+/syn-tdT+ colocalisation; cyan arrows indicate GFP+/syn-tdT+/VGLUT2+ colocalisation. Scale bars: 20 µm for all images except inset zoom in **b**: 10 µm. **e-l**, Quantification (mean ± SEM) of density (**e,i**), proportion glutamatergic (**f,j**), proportion within 5 µm of a HuC/D+ soma (**g,k**), and proportion contacting a VGLUT2^syn-tdT^ neuron (**h,l**) of GFP+/syn-tdT+ varicosities in the SI (**e-h**) and colon (**i-l**) across different regions of VGLUT2^Long^ and VGLUT2^Circ^ neurons. **m-q**, Representative images of VGLUT2^syn-tdT^ (red) neurons transduced by AAV-GFP (green) and immunolabelled for enkephalin (**m**), substance P (**n**), CCK (**o**), CGRP (**p**), and VIP (**q**). Scale bars: 5 µm. **r**, Heatmap showing proportion of VGLUT2^Long^ and VGLUT2^Circ^ neurons in the SI and colon that colocalise with neuropeptides shown in **m-q**. n = 2-7 mice, 3-6 neurons per mouse.

We next asked whether other neurotransmitters and neuropeptides are released from the same varicosities to communicate with both neurons and non-neuronal cell types. Enkephalin and the tachykinin substance P were both found in longitudinal and circumferential neuron varicosities in the SI and colon, though substance P showed a strong enrichment in the colon and specifically in VGLUT2^Circ^ (Fig. 3m,n,r). Interestingly, enkephalin was the only neuropeptide to be found in varicosities on both branches and the primary fibre (Fig. 3m). Cholecystokinin (CCK) and CGRP were only found in the branches of a minority of VGLUT2^Long^ in the SI and colon, while VIP was not found in any (Fig. 3o-r), despite previously observing colocalisation between Vip^tdT^ and VGLUT2^GFP^ (Fig. 1).

These data show that varicosities in VGLUT2 neurons are found primarily in the nests of VGLUT2^Circ^ neurons and the sparse branches of VGLUT2^Long^ neurons, where they are likely pre-synaptic sites to release glutamate and neuropeptides such as enkephalin and substance P to communicate with other neurons.

### Knocking out VGLUT2 in the ENS quickens total GI motility

Application of exogenous glutamate to intestinal preparations is known to be able to depolarise myenteric neurons and alter muscle contractility, but the necessity of glutamate in the ENS has not been previously tested. In part this is because of the difficulty in genetically knocking out glutamate or its transporters, as VGLUT2 homozygous knockout mice die shortly after birth^23^. To restrict VGLUT2 knockout to the ENS, we crossed VGLUT2-Flx mice with Wnt1-Cre, a neural crest marker known to be expressed in most ENS neurons and glia, but the offspring of this cross also died shortly after birth (data not shown). Instead, given the strong colocalisation between Penk and VGLUT2 in both longitudinal and circumferential neurons, we crossed VGLUT2-Flx with Penk-Cre to generate PenkCre-VGLUT2^flx/flx^ mutants and VGLUT2^flx/flx^ littermate controls. We reasoned that this would knock out VGLUT2 in the majority of glutamatergic ENS neurons, while still limiting potential non-ENS effects. PenkCre-VGLUT2^flx/flx^ offspring were viable and appeared healthy, albeit with more male offspring than female. PenkCre-VGLUT2^flx/flx^ males were significantly smaller than their control littermates, weighing 18.4 g compared to 24.2 g, while this difference was not as pronounced in females (Fig. 4a). Male colons were also slightly shorter, with a mean of 45.5 mm in mutants and 51.1 mm in controls, while no significant differences were observed for SI length (Fig. 4b,c).

**Fig. 4.**
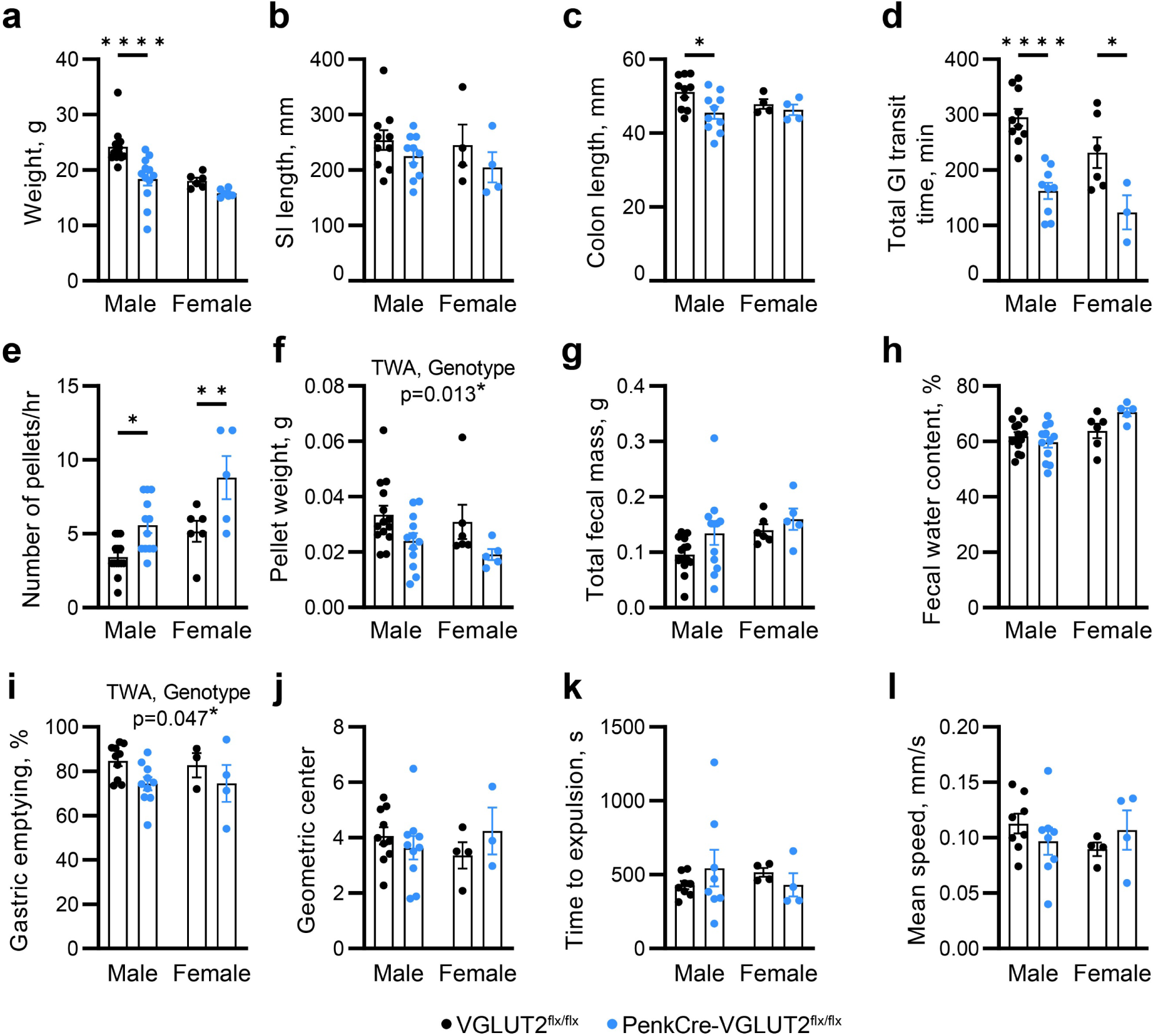
Knocking out VGLUT2 from Penk-Cre neurons accelerates total gastrointestinal transit. **a-c**, Weight, SI length and colon length of male and female PenkCre-VGLUT2^flx/flx^ mice (blue) and VGLUT2^flx/flx^ littermate controls (black). **d-h**, Total gastrointestinal transit time (**d**), number of pellets produced per hour (**e**), initial pellet weight (**f**), total faecal mass of all pellets (**g**), and faecal water content (**h**) following oral gavage of carmine red dye. Each dot for pellet weight and faecal water content represents the mean value of all pellets from a single mouse. Groups as in **a-c**. **i,j**, Percentage gastric emptying (**i**) and geometric centre of fluorescent signal (**j**) 15 minutes after oral gavage with rhodamine dextran. Groups as in **a-c**. **k,l,** Time to pellet expulsion (**k**) and mean pellet speed (**l**) of artificial faecal pellets inserted into the colon. Dots represent mean values from 4 trials per mouse colon. Groups as in **a-c**. n for **a-h**: VGLUT2^flx/flx^ male: 14; VGLUT2^flx/flx^ female: 6; PenkCre-VGLUT2^flx/flx^ male: 12; PenkCre-VGLUT2^flx/flx^ female: 5. n for **i-l**: VGLUT2^flx/flx^ male: 8-10; VGLUT2^flx/flx^ female: 4; PenkCre-VGLUT2^flx/flx^ male: 8-10; PenkCre-VGLUT2^flx/flx^ female: 4. All tests two-way ANOVAs (TWA) for genotype and sex. Comparisons in which there was an overall significant effect of genotype but no significant differences following multiple comparisons testing are indicated (**f,i**). *p< 0.05, **p< 0.01, ****p< 0.0001

To test *in vivo* GI function, mice were gavaged with carmine red, and the time elapsed until red faecal pellets were produced was measured. Mutants displayed significantly faster total GI transit time than littermate controls, with a mean of 163 and 124 minutes in male and female PenkCre-VGLUT2^flx/flx^ mice, respectively, compared with 295 and 230 minutes in their corresponding controls (Fig. 4d). This coincided with significantly more pellets produced by PenkCre-VGLUT2^flx/flx^ mice: 5.58 and 8.80 pellets/hr in males and females, respectively, compared with 3.43 and 5.17 pellets/hr in male and female controls. Both male and female PenkCre-VGLUT2^flx/flx^ pellets were significantly smaller than their respective controls’, weighing 28.55% and 38.13% less respectively, though the total faecal mass produced by each mouse and the faecal water content were not significantly different (Fig. 4e-h).

To determine the source of the differences in total GI transit within the GI tract, we separately tested stomach, SI and colon function. To test the stomach and SI, we gavaged mice with rhodamine dextran and waited for 15 minutes to observe gastric emptying and SI transit. There was a minor difference in gastric emptying between the groups, while there was no difference in SI transit (Fig. 4i,j). There was also no difference in colonic motility, assessed by the speed and transit of an artificial faecal pellet in *ex vivo* colons (Fig. 4k,l). Thus, while the source remains elusive, these data suggest that glutamatergic function in enkephalin neurons is important for *in vivo* functioning of the GI tract.

### Optogenetic activation of glutamatergic neurons stimulates colonic motility

To further explore the function of glutamate neurons within the ENS, we next asked whether optogenetic activation of enteric VGLUT2 neurons is sufficient to affect *ex vivo* colonic motility (Fig. 5a). We crossed VGLUT2-Cre with Cre-dependent channelrhodopsin2 to produce VGLUT2^ChR2^, and exposed a focal area (∼5 mm) of isolated colon to 460 nm LED light at 5 Hz for 20 seconds. Stimulation of the mid-colon of VGLUT2^ChR2^, but not ChR2 controls lacking Cre (WT^ChR2^), caused propulsion of an artificial faecal pellet through the colon and at greater speed compared with pre-stimulation conditions (Fig. 5b-d). This was neuronally mediated, as any response to optogenetic stimulation was abolished under tetrodotoxin (TTX; Extended Data Fig. 3a).

**Fig. 5.**
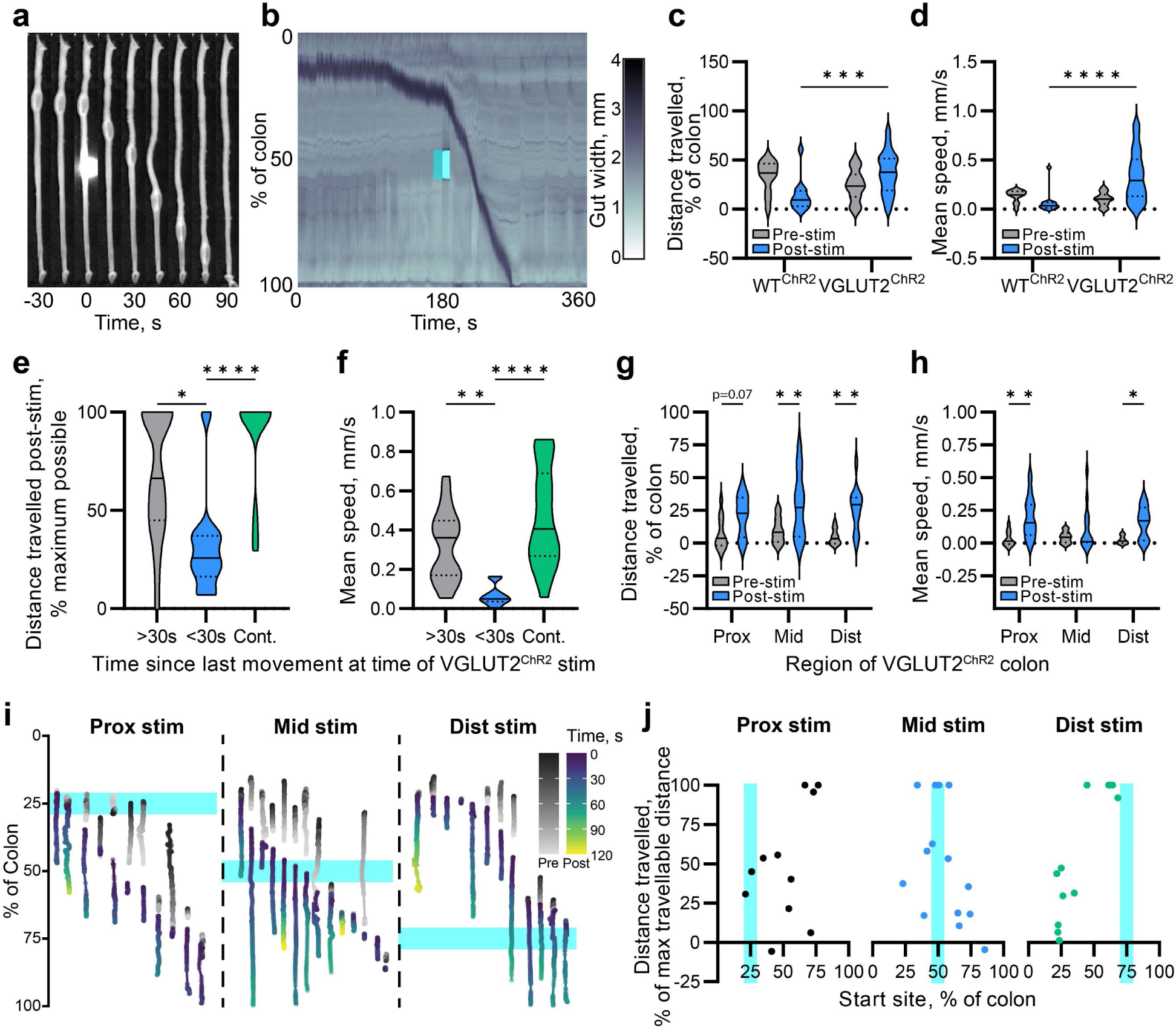
Optogenetic activation of VGLUT2 neurons stimulates colonic propulsive motility. **a,** Representative timelapse of an *ex vivo* VGLUT2^ChR2^ colon stimulated in the mid-colon with 460 nm LED at t=0. **b**, Representative spatio-temporal map of a VGLUT2^ChR2^ colon showing the width of each point along the length of the colon (y axis, %) over 6 minutes. Optogenetic stimulation (cyan box) occurs half way through the recording. The dark band indicates the artificial faecal pellet. **c,d**, Violin plots of distance travelled as a percentage of the full colon length (**c**) and mean speed (**d**) of artificial pellets before and after mid-colon optogenetic stimulation in WT^ChR2^ and VGLUT2^ChR2^ colons. n: WT^ChR2^: 6 mice, 2-3 trials per mouse; VGLUT2^ChR2^: 20 mice, 1-4 trials per mouse. **e,f**, Violin plots of the effect of time since last pellet movement on distance travelled by artificial pellets as a proportion of the remaining length of colon (**e**), and of mean pellet speed (**f**), following mid-colon optogenetic stimulation of VGLUT2^ChR2^ colons. n: >30s: 17 trials across 14 mice. <30s: 10 trials across 7 mice. Cont: 23 trials across 15 mice. 1-3 trials per mouse. **g,h**, Violin plots of the effect of optogenetic stimulation location on distance travelled as a percentage of the full colon length (**g**) and mean speed (**h**). n: Prox: 12 trials across 8 mice; Mid: 13 trials across 5 mice; Dist: 12 trials across 6 mice. **i**, Normalised motion tracks of individual artificial pellets in VGLUT2^ChR2^ colons, split based on stimulation location (cyan), coloured by time before (greys) and after (viridis) optogenetic stimulation. Movement in the x axis indicates colon displacement. Same dataset as **g,h**. **j**, Correlation between distance travelled by artificial pellets as a proportion of the remaining length of colon and the location of the pellet at the time of stimulation (start site), split based on stimulation location (cyan). Each dot represents a single trial. Same dataset as **g,h**.

Having established that VGLUT2 neuron activity could stimulate colonic motility, we next assessed factors that could affect the movement of the pellet through the colon in response to VGLUT2 neuron excitation. Factors included pellet location in the colon, stimulation location, and pre-stimulation state. Pre-stimulation activity had a strong effect on the subsequent influence of stimulation: if the colon had just finished moving the pellet, then stimulation was typically unable to elicit much response, whereas if the pellet was moving at the time of stimulation, it was highly likely that the pellet would be expelled (Fig. 5e,f). The location of stimulation also affected the response, albeit to a lesser degree, and, interestingly, exciting VGLUT2 neurons in any of the proximal, mid or distal colon was able to elicit a strong response in pellet propulsion, regardless of pellet location with respect to the stimulation location (Fig. 5g-j and Extended Data Fig. 3b,c). This was illustrated by the distal stimulation condition, which had a strong effect on pellet propulsion if the pellet was within the distal half of the colon, but could still affect proximally located pellets.

To determine if the effect of stimulating VGLUT2 cells was specific to VGLUT2 cells, we also stimulated Trpv1^ChR2^ colons, as Trpv1-Cre is expressed across a similar number of neurons to VGLUT2 (Extended Data Fig. 3d-f). Trpv1 neurons were primarily found within the MP, as well as some non-neuronal cells, most notably putative pericytes surrounding blood vessels within the SMP (Extended Data Fig. 3d-f). Exciting Trpv1 cells optogenetically had a minor effect on speed of propulsion, albeit less than stimulating VGLUT2 neurons, but did not have a corresponding effect on pellet distance travelled (Extended Data Fig. 3g,h).

These results demonstrate that VGLUT2 neurons are capable of strongly and specifically affecting motility across the length of the colon, but this can be modulated by stimulation location, luminal contents location, and the prior activity state of the enteric circuit.

### VGLUT2 neurons signal to a diverse array of different cell types

To establish how activation of VGLUT2 neurons achieves its effects on colonic motility, we sought to position VGLUT2 neurons within the enteric circuit by identifying which enteric neurons they communicated to. We focused our analysis on the respective branches and nests of VGLUT2^Long^ and VGLUT2^Circ^ neurons sparsely transduced by Cre-dependent AAV-GFP, as that was where syn-tdT signal and VGLUT2 IHC signal itself were most concentrated. We costained with 4 markers that marked the cell bodies of potential recipient enteric neuron subtypes: calretinin (excitatory motor neurons, ascending interneurons), nNOS (inhibitory motor), Scgn (putative interneuron and sensory), and Sst (putative interneurons, distinct from Scgn)(Fig. 6a-c).

**Fig. 6.**
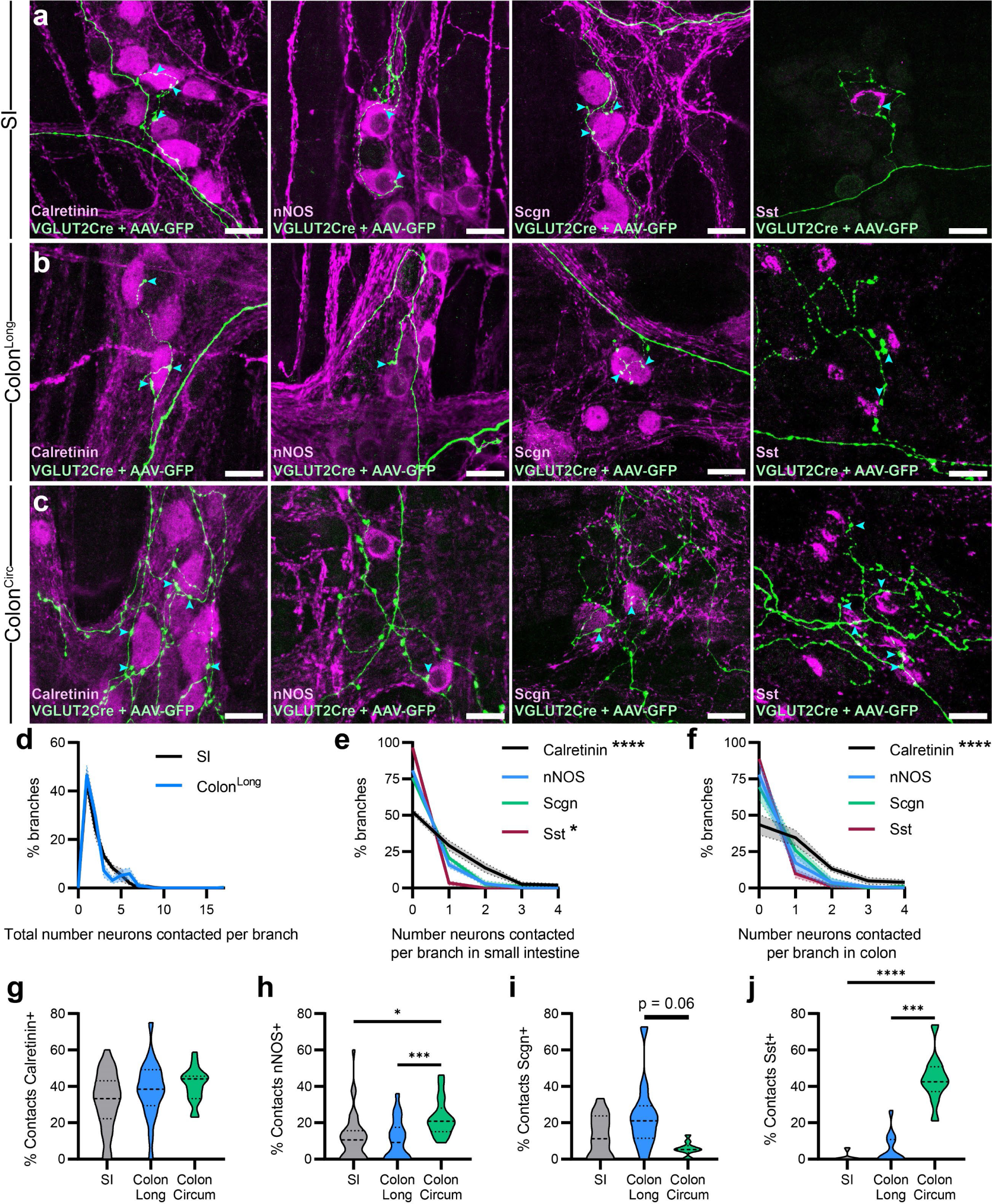
VGLUT2 neurons form putative synapses with diverse neuron populations. **a-c**, Representative images of adult wholemount MPs showing sparsely transduced VGLUT2^AAV-GFP^ fibres (green) from VGLUT2^Long^ neurons in the SI (**a**) and colon (**b**) and VGLUT2^Circ^ neurons (**c**), and immunohistochemical labels for calretinin, nNOS, Scgn and Sst (magenta). Cyan arrows indicate examples of VGLUT2 varicosities contacting indicated neuron type. Scale bars: 20 µm. **d**, Frequency distribution for the total number of HuC/D+ neurons contacted by VGLUT2^Long^ neuron branches. **e,f**, Frequency distribution showing number of neurons of a given subtype contacted by VGLUT2^Long^ neuron branches in the SI (**e**) and colon (**f**). Kruskal-Wallis test with Dunn’s multiple comparisons. n: ≥20 branches (VGLUT2^Long^) across 4-8 neurons per organ (SI or colon) per mouse, 6 mice. **g-j**, Proportion of total recipient neurons contacted by VGLUT2^Long^ and VGLUT2^Circ^ neurons positive for calretinin (**g**), nNOS (**h**), Scgn (**i**), or Sst (**j**). One-way ANOVA with Tukey’s multiple comparisons test. n: ≥20 branches (VGLUT2^Long^) or ≥5 nests (VGLUT2^Circ^) across 4-8 neurons per organ (SI or colon) per mouse, 2-4 mice per marker.

Approximately half of all VGLUT2^Long^ branches contacted only 1 neuron, and very few contacted more than 5 (Fig. 6d). Of the cell types investigated, Sst was the least likely to be contacted, while calretinin was the most likely, but all 4 cell types were in close contact with VGLUT2^AAV-GFP^ varicosities, in both the SI and colon (Fig. 6e,f). We next assessed the proportion of recipient neurons that were a given cell type out of the total recipient population, across VGLUT2^Long^ in the SI and both VGLUT2 classes in the colon (Fig. 6g-j). While no significant differences were found for calretinin and Scgn between VGLUT2^Circ^ and VGLUT2^Long^ neurons, VGLUT2^Circ^ neurons contacted approximately twice as many nNOS neurons and dramatically more Sst neurons as VGLUT2^Long^ did (Fig. 6g-j). Of note, no significant differences were found between SI and colon VGLUT2^Long^ neurons in their recipient populations. To determine if the proportion of recipient neurons that were a specific subtype was due to chance or due to preferential communication from VGLUT2 neurons, we performed chi-squared analysis, comparing the proportion of recipient neurons of a given cell type with the overall proportion of MP neurons of that cell type from an existing dataset^20^. SI VGLUT2^Long^ neurons did not show any preferential targeting, but did contact significantly fewer nNOS neurons than expected by chance (Extended Data Fig. 4a). Colonic VGLUT2^Long^ neurons preferentially targeted calretinin and Scgn neurons, and contacted fewer nNOS and Sst neurons than expected based on chance (Extended Data Fig. 4b). While VGLUT2^Circ^ neurons similarly showed a preference for calretinin neurons and avoidance of nNOS neurons, they contacted fewer Scgn neurons than expected by chance (Extended Data Fig. 4c). VGLUT2^Circ^ also preferentially targeted Sst neurons, with 43% of neurons receiving input from VGLUT2^Circ^ being Sst+, while Sst+ neurons represent only 13% of MP neurons in the proximal colon (Extended Data Fig. 4c). These data demonstrate that VGLUT2 neurons output to a diverse array of different cell types with different functions, and that VGLUT2^Long^ and VGLUT2^Circ^ neurons show distinct preferences in recipient populations.

### Calb1 and Prlr mark the two separate VGLUT2 populations in the colon

Given the strong differences in morphology and recipient neurons between VGLUT2^Long^ and VGLUT2^Circ^, we next asked whether the two populations of colonic neurons would have different contributions to the motility effect seen in optogenetics experiments (Fig. 5). Separate manipulation of the two morphological populations necessitated identifying single gene markers that could be used as Cre drivers. To identify appropriate single gene markers, we first matched the morphological classes to previously described transcriptional classes.

scRNAseq data predicted that there are three VGLUT2 neuron populations in the colon: putative interneuron 2 (PIN2), PIN3, and putative sensory neuron 4 (PSN4)^5^. Through manual interrogation of the dataset, we anticipated that the three populations could be distinguished by their expression of enkephalin, VIP, and secretagogin, where PIN3 would only express enkephalin, PSN4 would express VIP ± secretagogin, and PIN2 would express enkephalin ± secretagogin. VIP and enkephalin were expected to colocalize only a small amount in the PIN2 group; this colocalization was not tested directly.

To determine co-expression, we systemically injected Flp-dependent AAV-mCherry into mice expressing VGLUT2-Flp, Penk-Cre or Vip-Cre, and Cre-dependent ReaChR-mCitrine, creating _VGLUT2AAV-mCherry/PenkReaChR-mCitrine and VGLUT2AAV-mCherry/VipReaChR-mCitrine, respectively_ (Extended Data Fig. 5a-d). This would sparsely transduce VGLUT2 neurons to allow visualisation of morphology, and determine colocalisation with the mCitrine reporter in Penk or Vip neurons, and with Scgn IHC. Of 9 VGLUT2^Circ^ neurons in VGLUT2^AAV-mCherry^/Penk^ReaChR-mCitrine^ mice, all expressed mCitrine and very few expressed Scgn (2/9), which suggested that these were PIN2 or PIN3 neurons. However, we concluded that VGLUT2^Circ^ belonged to PIN3 because no VGLUT2^Circ^ neurons expressed mCitrine in VGLUT2^AAV-mCherry^/Vip^ReaChR-mCitrine^ mice (Extended Data Fig. 5a,b), as predicted by scRNAseq.

PIN2 and PSN4 were more difficult to separate, likely due to their similar transcriptional profiles^5^. Of 25 VGLUT2^Long^ neurons in VGLUT2^AAV-mCherry^/Penk^ReaChR-mCitrine^ mice, 22/25 expressed Penk, of which 8 expressed Scgn, which may suggest that these 22 neurons belonged to PIN2, with the remaining 3 Penk-neurons in PSN4 (Extended Data Fig. 5c). 10/13 VGLUT2^Long^ neurons in VGLUT2^AAV-mCherry^/Vip^ReaChR-mCitrine^ mice coexpressed mCitrine (Extended Data Fig. 5d). These VGLUT2+/VIP+ longitudinal neurons likely belong to both PIN2 and PSN4, and were harder to separate than expected due to the low number of Scgn-expressing neurons (3/10). It is important to note that no differences in morphology between the putative PIN2 and PSN4 longitudinal neurons were observed.

Having established that VGLUT2^Circ^ neurons belong to PIN3 and VGLUT2^Long^ belong to PIN2/PSN4, we interrogated scRNAseq datasets to identify single gene markers that were highly enriched in each population but with minimal expression elsewhere in the ENS. Consequently, PIN3 (VGLUT2^Circ^) was marked by prolactin receptor (*Prlr*), which shows widespread expression in both males and females throughout the body, including neurons and non-neurons in the intestines^24^ (Extended Data Fig. 5e). In contrast, PIN2 (VGLUT2^Long^) was marked by calbindin (*Calb1*), a calcium binding protein that has previously been used as a sensory marker^1^, though it has also been posited as an interneuron marker^25^. PIN2 was chosen over PSN4 to represent VGLUT2^Long^ neurons because PIN2 had more readily available unique markers to distinguish it from other enteric neuron populations (including PSN4), such as *Calb1*, *Piezo2*, and *Bdnf*.

We validated co-expression of VGLUT2 with both candidates using combined RNAscope/IHC for *Slc17a6, Prlr*, *Calb1* and HuC/D (Extended Data Fig. 5f), though cell counts of *Slc17a6*+/*Prlr*+ cells proved difficult due to the low number of transcripts per neuron. Quantification of *Calb1*+ expression in VGLUT2 neurons was performed using combined RNAscope for *Calb1* and IHC for VGLUT2^tdT^ and HuC/D. *Calb1* coexpression showed significant regional variation, being highest (22% of VGLUT2 neurons) in the proximal colon, and lowest (8%) in the mid-colon (MC) (Fig. 7a,b).

**Fig. 7.**
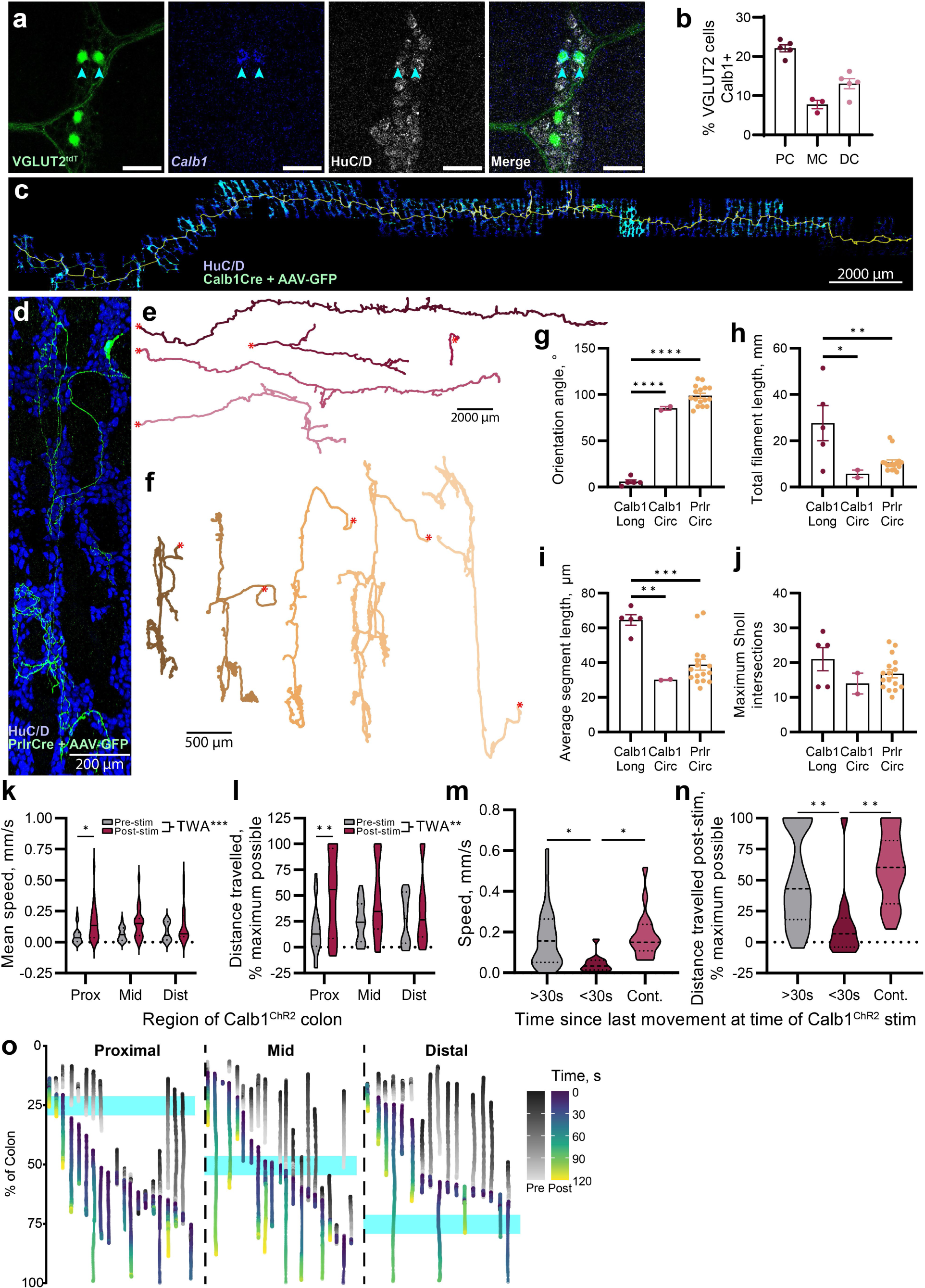
VGLUT2 neuron classes can be isolated genetically. **a**, Representative images of adult wholemount MP showing IHC for VGLUT2^tdT^ (green) and HuC/D (grey) alongside *Calb1* RNAscope (blue). Cyan arrows indicate VGLUT2+/*Calb1*+ neurons. Scale bars: 50 µm. **b**, Proportion of VGLUT2 neurons (mean ± SEM) positive for *Calb1* in the MP across colonic regions. **c,d**, Representative images of Calb1-Cre neurons (**c**) and Prlr-Cre (**d**) transduced by Cre-dependent AAV-GFP and immunostained for GFP (green) to reveal full neuron morphology, alongside HuC/D (blue). Scale bars as indicated. **e,f**, Representative traces of AAV-GFP-labelled Calb1-Cre (**e**) and Prlr-Cre (**f**) neurons in the colon. Soma location indicated by red asterisk. Scale bars as indicated. **g-j**, Quantification (mean ± SEM) of Calb1^AAV-GFP^ and Prlr^AAV-GFP^ neuron orientation (**g**), total filament length (**h**), average segment length (**i**), and maximum Sholl intersections (**j**). n: Calb1^Long^: 5, Calb1^Circ^: 2, Prlr: 16. Each dot represents a different neuron. **k,l**, Violin plots of the effect of optogenetic stimulation location on mean speed (**k**) and distance travelled as a percentage of the remaining colon length (**l**) in Calb1^ChR2^ colons. Two-way ANOVA (TWA) for pre/post stimulation and stimulation location. Significant differences were observed for overall pre-vs post-stimulation, and Šídák’s multiple comparisons test revealed a significant difference between proximally stimulated samples. n: 17-21 trials over 8 mice per region stimulated. **m,n**, Violin plots of the effect of time since last pellet movement on mean pellet speed (**m**) and distance travelled by artificial pellets as a proportion of the remaining length of colon (**n**) following mid-colon optogenetic stimulation of Calb1^ChR2^ colons. n: >30s: 29 trials across 8 mice. <30s: 12 trials across 6 mice. Cont: 16 trials across 6 mice. 1-6 trials per mouse. **o**, Normalised motion tracks of individual artificial pellets in Calb1^ChR2^ colons, split based on stimulation location (cyan), coloured by time before (greys) and after (viridis) optogenetic stimulation. Movement in the x axis indicates colon displacement. Same dataset as **k-n**.

We next confirmed the respective longitudinal and circumferential morphology of Calb1 and Prlr neurons using specific Cre lines^26^ and sparse viral transduction (Fig. 7c-f). For Calb1 neurons, we found no evidence of epithelial projections, as might be expected of sensory neurons, with the majority of Calb1 neurons showing descending longitudinal interneuron morphology, while 2 Calb1 neurons showed circumferential morphology (Fig. 7c,e,g-j). 100% of Prlr neurons analysed showed circumferential morphology closely resembling VGLUT2^Circ^ (Fig. 7d,f-j).

We focused on the function of Calb1 neurons using the same *ex vivo* optogenetics approach as before (Fig. 5). This approach was not feasible with the Prlr population, as crossing Prlr-IRES-Cre with a CAG-controlled ChR2 line would likely lead to off-target, non-neuronal effects following stimulation given the widespread expression of Prlr. Stimulating Calb1^ChR2^ neurons had a similar effect to stimulating VGLUT2^ChR2^, increasing the speed and distance travelled of artificial pellets (Fig. 7k,l). These effects also showed the same dependency on pre-stimulation state (Fig. 7m,n), and activated Calb1^ChR2^ neurons were able to stimulate organ-wide motility regardless of stimulation site (Fig. 7o, Extended Data Fig. 5g). Thus, it is likely that VGLUT2^Long^ neurons in the colon, marked by Calb1, are responsible for the ability of VGLUT2^ChR2^ to initiate propulsive motility in the colon.

## Discussion

We present a thorough morphological and functional characterization of glutamatergic circuitry within the enteric nervous system. VGLUT2 neurons present as putative longitudinal interneurons, projecting over long distances in the descending direction within the myenteric plexus of the small intestine and colon. Additionally, we uncover that VGLUT2 is expressed in a novel class of circumferential neurons in the colon, a previously unrecognized component of the enteric network. VGLUT2^Long^ neurons primarily establish glutamatergic synapses at branches and terminals, while VGLUT2^Circ^ neurons form synapses in nests, extensively innervating specific myenteric ganglia. Knocking out glutamate from enkephalin neurons considerably accelerates total GI transit and affects faecal pellet production. Furthermore, stimulation of VGLUT2 neurons *ex vivo* initiates strong propulsive motility, likely facilitated by Calb1+ longitudinal glutamatergic neurons, engaging multiple components of the enteric circuit by directly communicating with a variety of distinct neuronal subtypes.

Our sparse viral transduction demonstrates that VGLUT2^Long^ neuron fibres never leave the plane of the MP, showing no direct interaction with other intestinal layers and suggesting an interneuron identity. This classification is further supported by co-expression with 5-HT, VIP and enkephalin^3,27,28^. This contrasts with prior speculation that enteric glutamatergic neurons are sensory, on the basis of co-expression with substance P and calbindin^9^. While we note similar co-expression, substance P is found across a wide variety of neuron subtypes^6^, and our tracing of calbindin neurons does not suggest a classic sensory morphology, given the lack of projections to the epithelium, which we did observe for VIP neurons. Glutamate neurons have also previously been noted in the SMP, which our data do not support^9^. This discrepancy may be due to the previous study employing IHC of glutamate directly and therefore potentially identifying GABA neurons in which glutamate is a precursor^29^.

In contrast to VGLUT2^Long^, the function of VGLUT2^Circ^ neurons, to our knowledge a novel class of enteric neuron, remains unclear. PrlrCre is an effective tool for isolating VGLUT2^Circ^ neurons, but because of its widespread non-neuronal expression in the intestines, we could not use PrlrCre with ChR2 or other lines that were not neuronally restricted. Though circumferentially oriented neurons have been identified previously and suggested to be sensory^22^, VGLUT2^Circ^ neurons are distinct from these by being monoaxonal and less arborised. VGLUT2^Circ^ neurons tend to innervate one stripe of neurons in the MP, suggesting that they could be involved in coordinating circumferentially aligned ganglia^20^.

We found that VGLUT2 neurons were able to strongly and swiftly initiate propulsion of luminal contents following *ex vivo* optogenetic stimulation. VGLUT2 is also expressed in some extrinsic, non-ENS fibres within the intestinal wall, particularly in dorsal root ganglion sensory fibres, which can release CGRP from their sensory terminals to affect GI motility through inflammation and sensitisation to pain^30^. These terminals, which may still be present in the *ex vivo* preparation, also express Trpv1, thus because we saw only a minor effect on pellet propulsion when stimulating Trpv1 neurons, it is unlikely that these terminals are responsible for the effect seen when stimulating VGLUT2 neurons. As shown by our synaptic tracing, motility initiation and acceleration instead appear to be achieved by VGLUT2 neurons engaging multiple components of the enteric circuit. These include other VGLUT2 neurons, supporting the idea of interneuronal chains for long distance communication beyond local microcircuits^31,32^, and calretinin neurons. Calretinin neurons represent both motor neurons, which directly control motility, and ascending interneurons, which could allow signalling oral to the site of stimulation^33^. High speed and large field-of-view calcium imaging following stimulation of specific neurons would be necessary to fully visualise the flow of information through the enteric circuit.

Though confocal microscopy cannot be used to definitively identify synaptic contacts, the approach has been validated by electron microscopy and used to establish enteric circuitry previously^11,34,35^, and the preferential targeting by VGLUT2 neurons to different neuronal subtypes in the recipient population suggests specificity. The majority of synaptic sites on VGLUT2^Long^ neurons are on branches, but they are also present on the primary fibre, particularly at the terminal, where they often do not contact HuC/D+ cells. It is possible that synapses are formed between an axon and unlabelled dendrite distal to the soma, rather than between axon and soma, and indeed commonplace in the central nervous system. Dendrites in the ENS receive the majority of synaptic input, but are typically small, filamentous, and very close to the soma^36–38^, with only circumferential neurons displaying anything resembling longer dendrites amongst the neuron types we imaged. This may suggest that putative synaptic sites away from other neurons are instead involved in communicating with non-neuronal cell types, or in axo-axonal communication. Support for this includes the observation that enkephalin was found in varicosities on the primary fibre, which has recently been proposed as a signalling molecule between enteric interneurons and non-neurons such as colonocytes and T cells, based on receptor-ligand pair mapping^5^.

While the cell-type specific KO and optogenetic experiments provide strong evidence for the role of glutamatergic neurons in GI motility, the molecular role of glutamate in achieving these effects remains to be elucidated. Glutamate release from varicosities in colonic motility have previously been suggested to induce and alter the strength of muscle contractions via ionotropic receptors, mediate slow synaptic transmission via metabotropic glutamate receptors, and facilitate synaptic plasticity in the ENS, highlighting the complexity of glutamatergic neurotransmission^7–11,19^. Given the plethora of different glutamatergic receptors present on enteric neurons, it is likely that glutamate has multiple functional roles, depending on circuit state and whether it is released from VGLUT2^Long^ or VGLUT2^Circ^. Glutamate release from a source that is extrinsic to the ENS may also have played a role in disrupting GI transit in the cell-type specific KO experiments, due to Penk/VGLUT2 co-expression in areas of the hindbrain and spinal cord, both of which can modulate GI motility^17,39–42^. Finally, optogenetic activation will result in release not only of glutamate but of co-expressed signalling molecules as well, including ACh, enkephalin and substance P, all of which may modulate circuit activity in different manners. Future experiments should seek to disentangle the effects of glutamate from co-released molecules.

In conclusion, our findings demonstrate that intestinal motility is regulated by glutamatergic neurons, which in turn can be segregated into at least two distinct subtypes, one of which represents a novel neuron class in the ENS. Our studies represent a step forward in elucidating the complexity of enteric circuits, demonstrating that defined neuronal subtypes communicate directly with a wide variety of other neuron subtypes to facilitate long-distance communication in the intestines beyond immediate sensory responses to local luminal contents. Future work should further explore the active roles that interneurons play in processing information in the ENS and in facilitating plasticity in response to physiological and disease states.

## Materials and Methods

### Animals

All procedures conformed to the National Institutes of Health Guidelines for the Care and Use of Laboratory Animals and were approved by the Stanford University Administrative Panel on Laboratory Animal Care. Mice were group housed up to a maximum of five adults per cage. Food and water were provided *ad libitum* and mice were maintained on a 12:12 LD cycle. All experiments were performed on adult mice aged 2-10 months of both sexes.

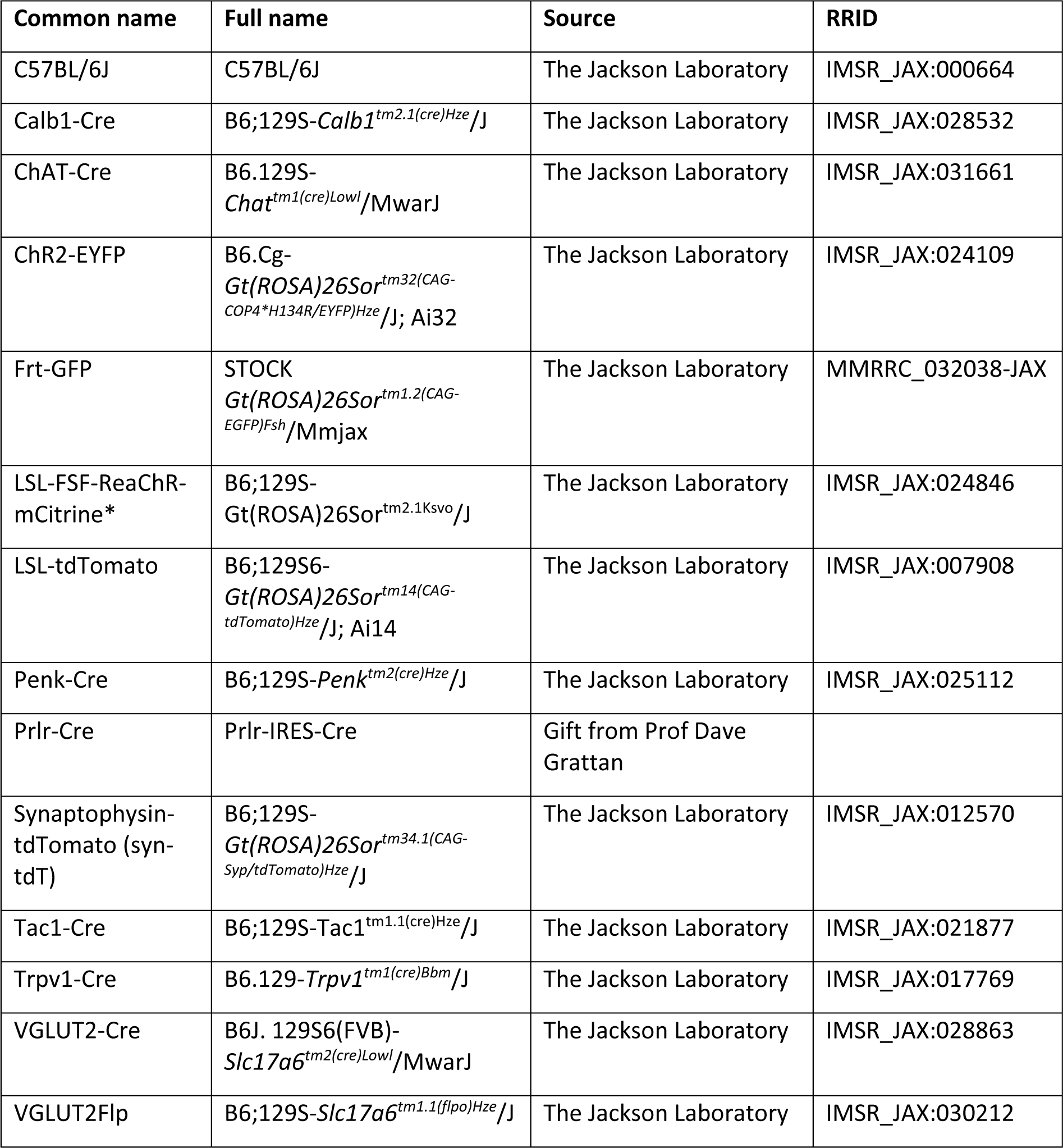
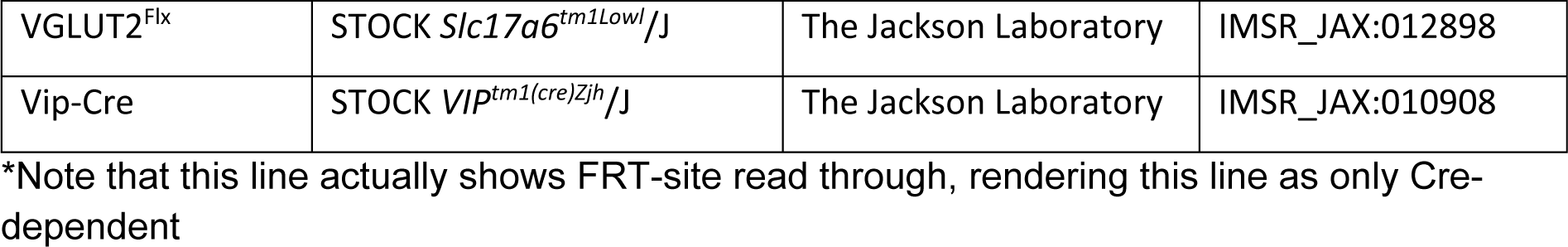

### Immunohistochemistry

Immunohistochemistry was performed as previously described^20^. Mice were euthanized by CO_2_ and cervical dislocation and the intestines were removed and flushed of faecal contents with cold PBS. The proximal, middle and distal 2-3 cm of the small intestine were separated to isolate the duodenum, jejunum and ileum, respectively, while the colon was cut in two. All segments were pinned to Sylgard 170 in cold PBS, and the mesentery was cut away before cutting open each segment longitudinally along the mesenteric border. Each piece of tissue was stretched flat and pinned, muscularis facing upwards, for fixation in 4% PFA for 90 minutes at 4°C with shaking. After fixation, the muscularis was peeled away using fine forceps and a cotton bud.

For immunohistochemistry, small pieces of wholemount tissue (typically ∼7×7 mm^2^) were placed in WHO microtitration trays (International Scientific Supplies) containing PBS. Tissue was incubated in PBT (PBS, 1% BSA, 0.1% Triton X-100) containing the primary antibodies overnight at 4°C with shaking. The following day, tissue was washed 3 times in PBT for 30 minutes each. Tissue was then transferred to PBT containing secondary antibodies, incubated for 2 h at room temperature with shaking. After washing in PBT and PBS, tissue was mounted onto slides using a paintbrush, ensuring it was flat by gentle manipulation with paint brushes under a dissection microscope. The tissue was rinsed in ddH2O after air-drying and coverslipped using Fluoromount-G (Southern Biotech).

### RNAscope

Dissections and tissue processing for RNAscope were performed as described for immunohistochemistry, but after 90-minute fixation and peeling, they were returned to fresh 4% PFA and further fixed at 4°C overnight. Protein-RNA co-detection was performed as previously described^44^ using RNA-protein Co-detection ancillary kit (ACD 323180), adapted for wholemount tissue. Tissue was placed in staining nets and dehydrated in ethanol (50%, 70%, 100%, 100%) for 5 minutes each before hydrogen peroxide incubation for 15 minutes. After a brief rinse in water, tissue was incubated in co-detect target antigen retrieval solution for 5 minutes in a steamer at >95°C, then rinsed in PBS and incubated overnight with primary antibody (Table 1) diluted in co-detection diluent. RNA detection was then performed using RNAscope multiplex fluorescent reagent kit V2 (ACD 323100). Tissue was washed in PBS with 0.2% Tween (PBS-T), and post-fixed in 10% formalin for 30 minutes. After further PBS-T washes, tissue was digested with Protease Plus for 30 minutes at 40°C. Following a rinse with water, tissue was incubated for 2 hours in RNAscope probes (Table 3) at 40°C. Amplification and development of probe signal was performed according to manufacturer’s instructions. After probe development, tissue was incubated with appropriate secondary antibodies (Table 2), then mounted and coverslipped on slides as for IHC.

**Table 1.**
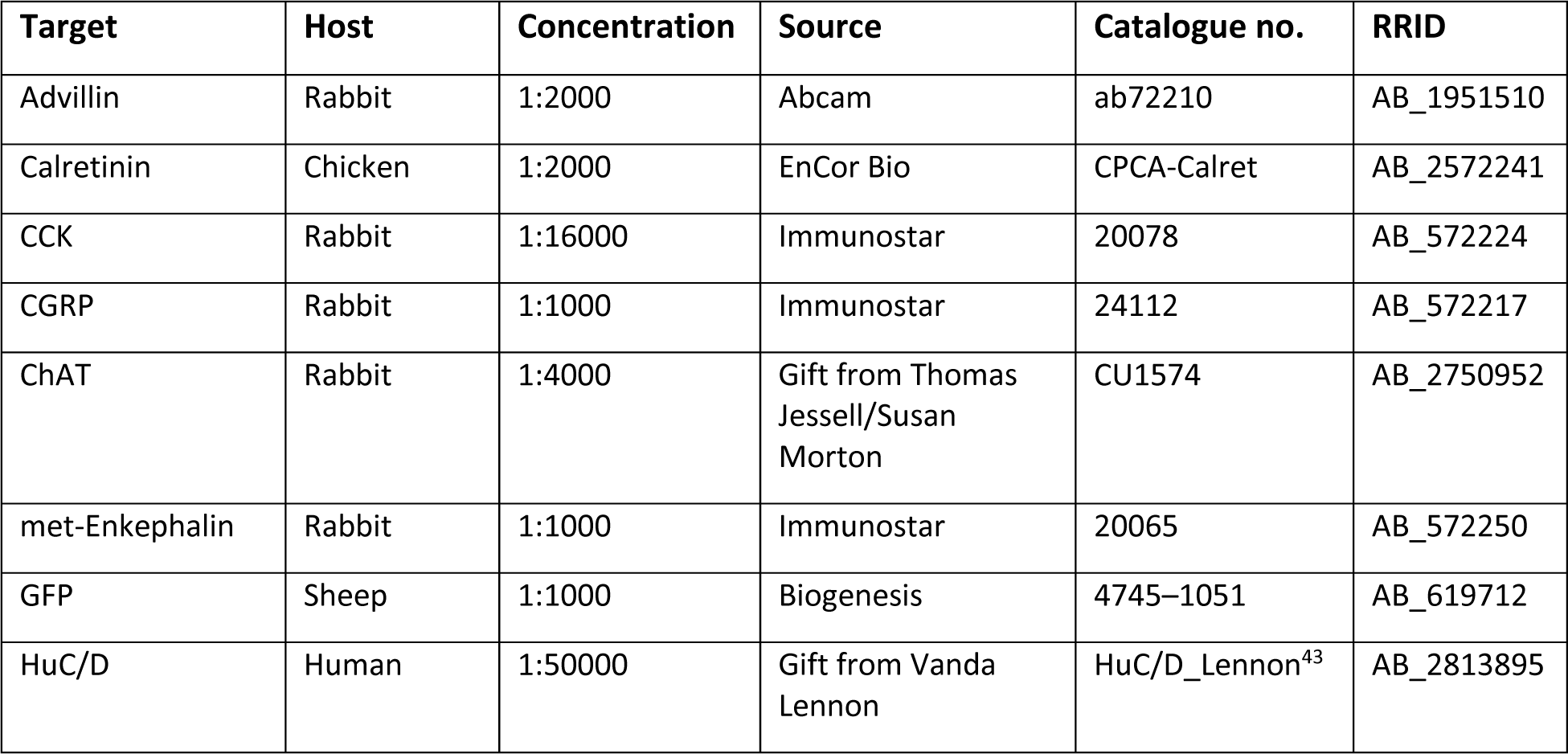
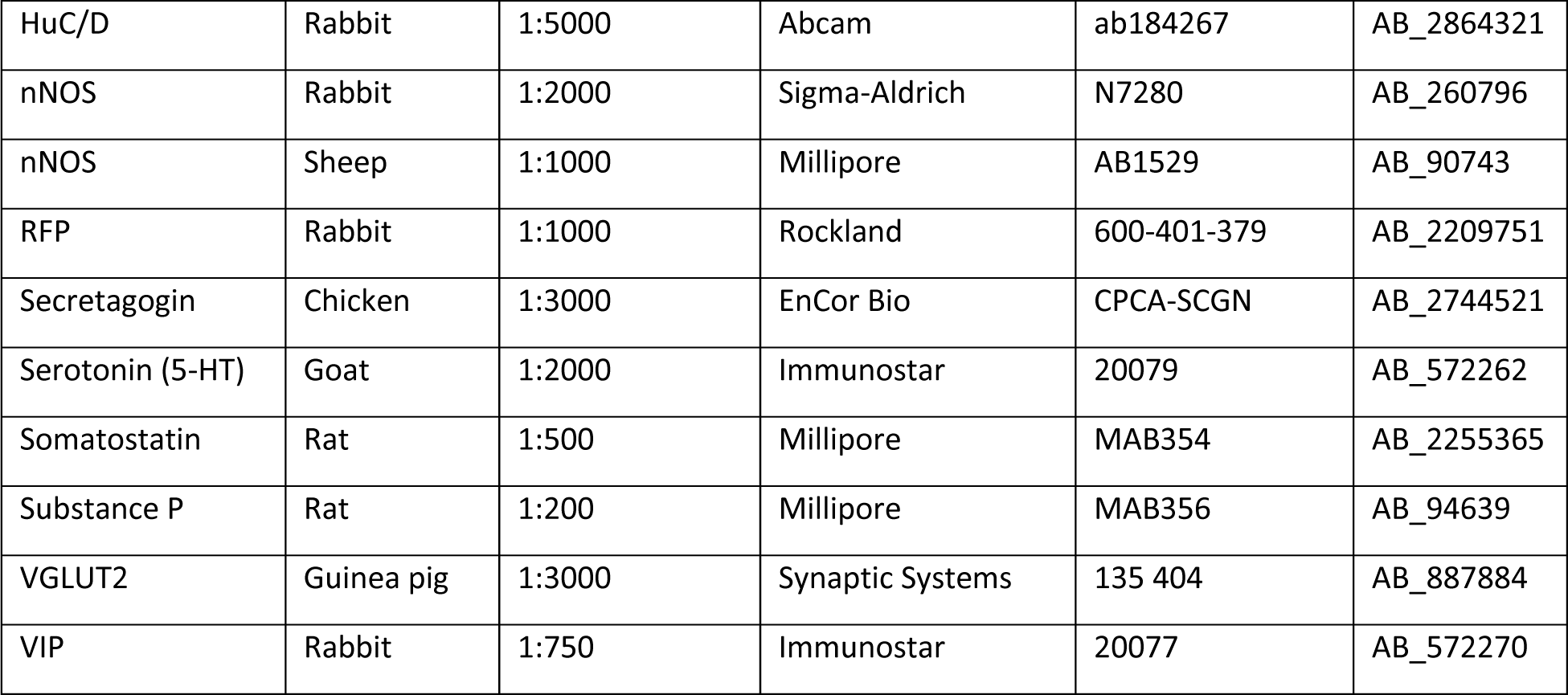
Primary antibodies.

**Table 2.**
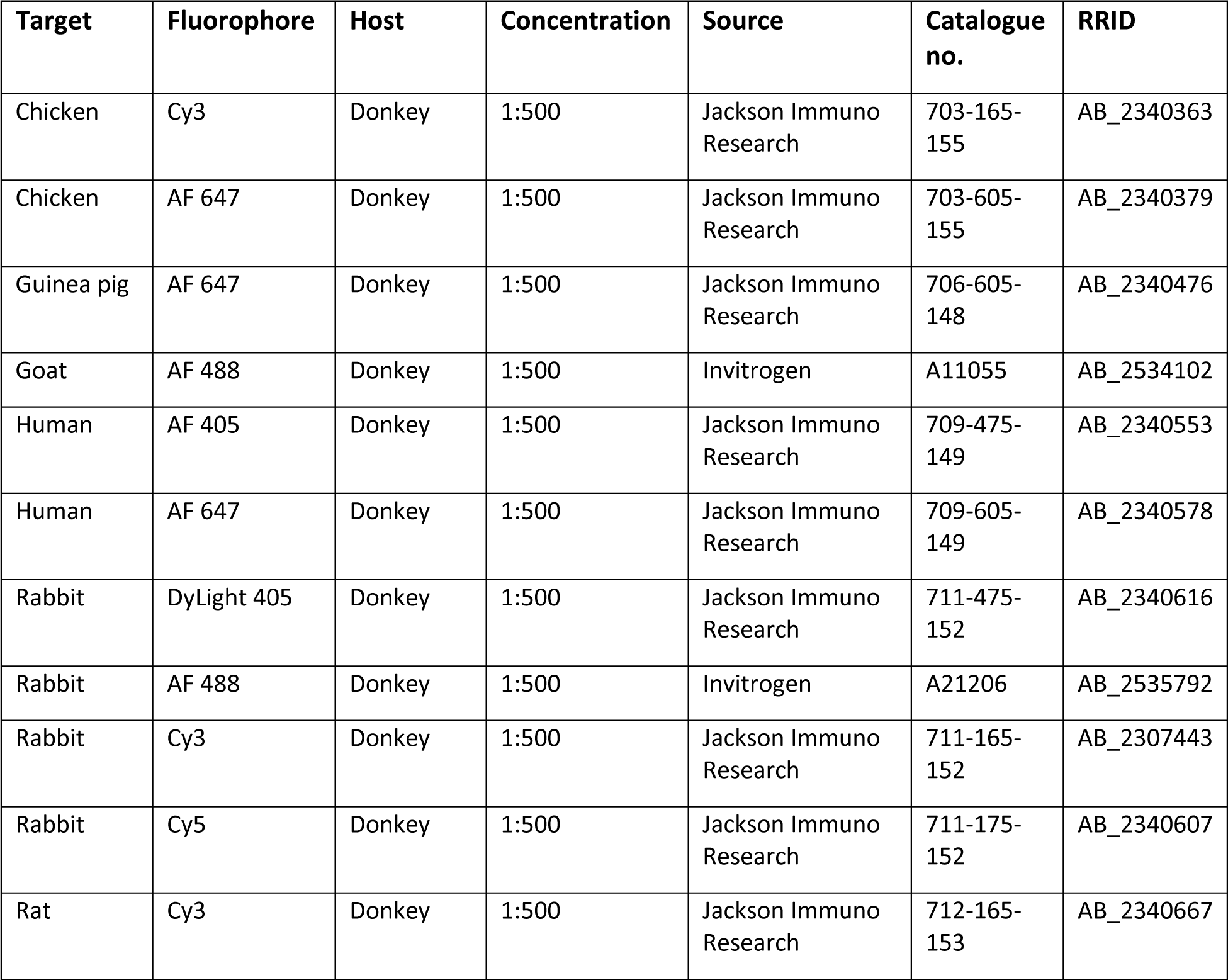
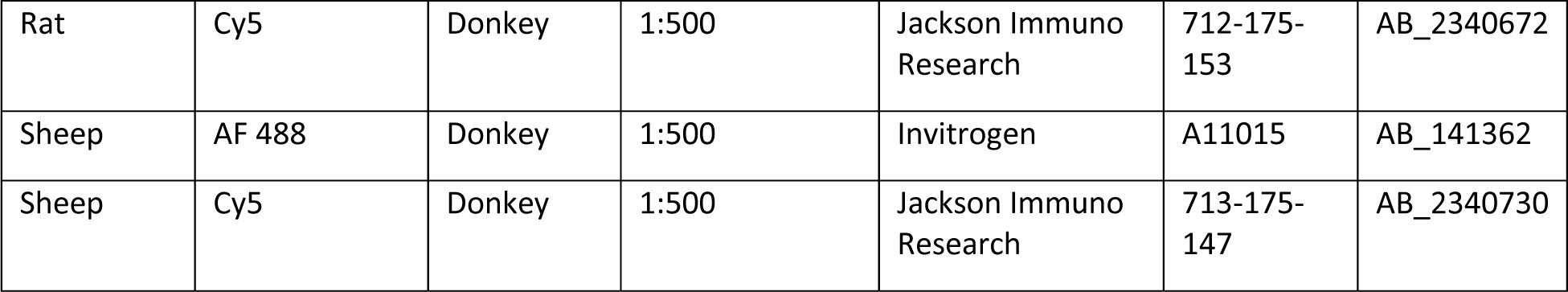
Secondary antibodies.

**Table 3.**
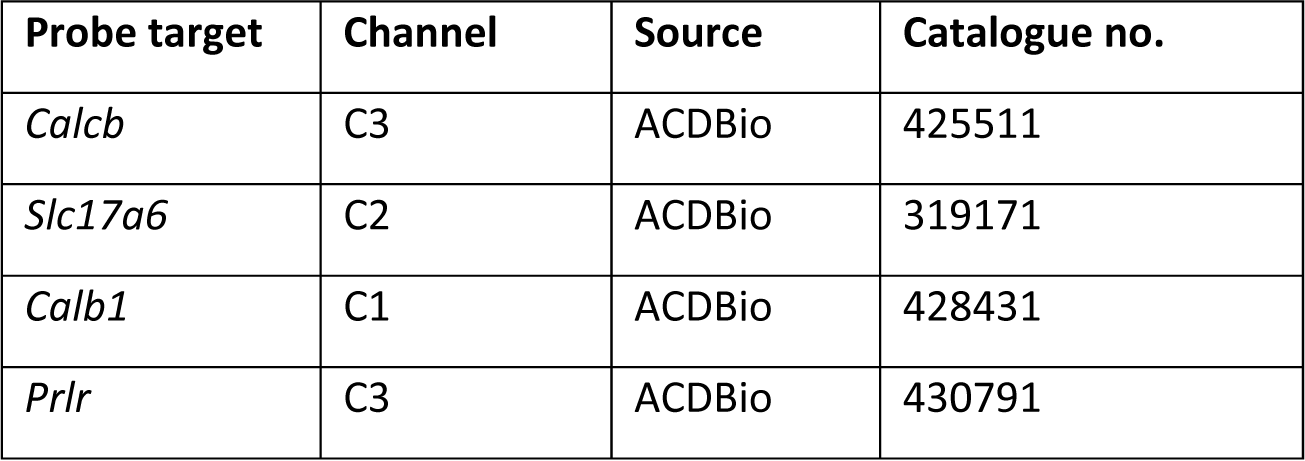
RNAscope probes.

| Probe target | Channel | Source | Catalogue no. |
| --- | --- | --- | --- |
| *Calcb* | C3 | ACDBio | 425511 |
| *Slc17a6* | C2 | ACDBio | 319171 |
| *Calb1* | C1 | ACDBio | 428431 |
| *Prlr* | C3 | ACDBio | 430791 |

### *Ex vivo* colonic motility

*Ex vivo* colonic motility was assessed using an experimental setup modified from approaches previously described^45,46^. The colon was dissected out from 2-10 month-old mice euthanized by CO_2_ and cervical dislocation, and submerged in warmed, carbogenated Krebs solution (pH 7.4 containing (in mmol/l): 117 NaCl, 4.7 KCl, 3.3 CaCl_2_, 1.5 MgCl_2_, 25 NaHCO_3_, 1.2 NaH_2_PO_4_ and 11 Glucose) in a Sylgard dish. For experiments using an artificial faecal pellet, the caecum was removed at this point and endogenous faecal matter was gently flushed from the colon using a syringe and warmed Krebs solution; otherwise, the caecum and faecal matter were left intact. The mesentery was cut away and the colon was transferred to an organ chamber, which was continuously perfused with carbogenated warm Krebs and heated from below by a water bath to maintain the chamber at ∼35°C. The colon was pinned at either end to the Sylgard chamber base under light tension. After 10 minutes’ acclimation, videos were recorded at 3.75 frames/s using IC capture software (Imaging Source) and a high-resolution monochromatic firewire industrial camera (Imaging Source, DMK41AF02) connected to a 2/3″ 16mmf/1.4 C-Mount Fixed Focal Lens (Fujinon HF16SA1) mounted above the organ bath. If used, tetrodotoxin (TTX, 1 µM; Alomone Labs) was diluted in warm, carbogenated Krebs and exchanged with the organ bath buffer, then circulated.

For experiments involving artificial pellets described below, 3D-printed pellets based on real faecal pellets were lubricated with KY jelly, inserted into the proximal colon and gently pushed ∼1.5 cm in using a blunt-ended gavage needle.

Experiments assessing colonic motility of PenkCre-VGLUT2^flx/flx^ mice were performed at the same time as SI transit and gastric emptying experiments (see below), thus mice were fasted overnight prior to dissection. Artificial pellets were Clear Resin v4 (Formlabs) and were based on real faecal pellet sizes from the experimental mice, given that PenkCre-VGLUT2^flx/flx^ mice produced smaller pellets and had smaller colons. Pellet widths were 2.4 mm and 3 mm for mutants and controls, respectively. Experimental trials were defined as a single passage of the artificial pellet through the colon to expulsion. Four trials were run per colon, with the artificial pellet being re-lubricated and re-inserted each time it was expelled.

For optogenetics experiments, 6-minute videos were acquired in which the colon was stimulated during the midpoint of the video by a 460 nm LED (UHP-T-460-DI, Prizmatix; 20 s stimulation, 5 Hz, 20 ms pulse-width^47^, controlled using a signal generator (Feeltech FY6600-60M)) situated 25%, 50% or 75% along the length of the colon for proximal, mid or distal stimulation, respectively. Artificial pellets were 3D-printed polycarbonate of approximately 5 mm in length and 3 mm at their widest. The location of the pellet was not controlled by the experimenter, and was based on its location at the time of stimulation. So that the fibre optic and associated apparatus did not interfere with the video recording, the light was emitted ∼20 mm away from the colon from a pinhole in the centre of a shield placed over the end of a collimator (Prizmatix), resulting in an illumination spot on the colon of ∼5 mm diameter. Experimental trials involving stimulation began only once the pellet had been inserted and expelled by the colon in the absence of any stimulation, to confirm normal colonic activity. The pellet was then re-inserted for experimental trials involving stimulation.

### *In vivo* procedures

#### Total GI transit

Whole GI transit was assessed as previously described^46^. Mice were orally gavaged with 300 µl 6% Carmine red (C1022; Sigma-Aldrich) in 0.5% methylcellulose (274429; Sigma-Aldrich) dissolved in 0.9% NaCl. Mice were separated into individual cages containing only a cotton nestlet square and a small weigh boat containing gel (DietGel 76A), then observed for up to 7 hours for production of red faecal pellets.

#### Faecal water content, pellet number and pellet size

Faecal water content was assessed as previously described^46^. Mice were housed individually in empty cages for 1 hour during which all faecal pellets were collected immediately after expulsion, photographed, and stored in pre-weighed tubes. Tubes were weighed at the end of the 1-hour observation to determine faecal mass, then incubated for 48 hours at 55°C. The dried pellets were then weighed and compared to their original faecal mass to determine faecal water content. Pellet length was measured using FIJI.

#### Small intestine transit and gastric emptying

SI transit and gastric emptying were determined as previously described^46^. Mice were fasted overnight prior to this experiment. Mice were orally gavaged with 100 µl 2.5 mg/ml rhodamine B dextran (D-1841; Molecular Probes) in 2% methylcellulose dissolved in water. 15 minutes after gavage, the mice were euthanized and the stomach and SI were removed in warmed, carbogenated Krebs buffer. The SI was measured and divided into 10 equal segments; the stomach and each SI segment were placed in separate tubes containing 0.9% NaCl and homogenized to release luminal contents. Following a 15-minute centrifugation at 2234 rcf, the fluorescence intensity of the supernatant was measured for each sample using a Varioskan LUX (Thermo Scientific).

### Retro-orbital injections

AAVs (AAV9::*CAG-FLEX-EGFP-WPRE*, Addgene 51502; AAV9::*Ef1a-fDiO-mCherry*, Addgene 114471) were diluted in sterile, ice-cold PBS to between 4 x 10^10^ and 4 x 10^11^ genome copies(GC)/ml, depending on the experimental aim and the Cre line being injected; injected AAV was diluted more for Cre lines with high representation in the ENS and for full neuron morphology tracing, in which sparse labelling was essential. Mice were anaesthetized with 3% isoflurane and treated with proparacaine droplets applied to the eye. After allowing time (30-60 s) for the proparacaine to take effect, mice were injected retro-orbitally with 100 μl diluted AAV using 31g insulin needles (BD-324920; BD). Mice were dissected after 2-4 weeks, allowing for adequate AAV expression.

### Image acquisition

Images were acquired using a 20x (NA 0.75) or 63x (NA 1.40) oil objective on a Leica SP8 confocal microscope. Regions to be imaged, including entire neurons for morphology analysis, were identified, acquired and stitched using the Navigator mode within LASX (Leica). For neuron morphology tracing, a preview of the entire neuron was built up in Navigator mode before acquiring a Z-stack of the entire region to ensure that all parts of the neuron in all layers (e.g. in muscle layers or MP) were included. Stacks were acquired with 2-2.5 μm between each focal plane. Branch or axon end points were identified by the abrupt loss of fluorescence signal in the fibre; neurons in which fluorescence signal gradually faded away were not imaged as fibre terminals could not be confirmed. Neurons were only included for morphology tracing if fibres could be confidently assigned to a given neuron.

### Image analysis

#### Cell counting

Cell counting analysis was performed as previously described^20^ using ImageJ/FIJI (NIH, Bethesda, MD). To count individual cell types, such as VGLUT2 neurons or VGLUT2 subtypes, first, Z-stacks of HuC/D were blurred using a Gaussian blur before thresholding and watershedding to identify individual neurons; neuronal locations were then identified and drawn using the Analyze Particles function (minimum size 50 µm^2^, minimum circularity 0.3), converting the output to a binary mask. The HuC/D mask was then combined with non-thresholded cell subtype images using the Image Calculator function. The result of this calculation was then maximally projected and counted in an automated fashion using the same procedure as described for HuC/D. For VGLUT2 overlap analysis, once VGLUT2 neuron locations were identified, they were combined with images of other markers (e.g. PenkCre-tdT, Scgn etc.) using Image Calculator, thresholded and counted as above to determine proportion of VGLUT2 neurons coexpressing these markers.

#### Neuron morphology

Three-dimensional reconstructions of neuronal morphology were created and analyzed in Imaris 9.7 (Bitplane). Fibres were traced in a semi-automatic way using the Autopath feature within the FilamentTracer module. Starting points were manually selected, typically at the neuronal soma. Fibre width in Autopath was set at 0.9 µm for tracing to enable tracing of fibres in close proximity. In cases of ambiguity, such as overlapping or recursive branches, the most parsimonious option was chosen (e.g. fewest number of branch points), ensuring that fibres never looped and reconnected with themselves. Filament statistics, including Sholl analysis and filament length, were exported to Microsoft Excel for grouping and further analysis. Assignment of neuron morphology to different functional groups was based on the following criteria. Interneurons were assigned if the neuron fibre stayed within the plane of the MP for all or the vast majority of its projection, and terminated there; ascending if they projected orally, descending if they projected aborally. Motor neurons were assigned if arborization within the muscle layers was observed; excitatory if they projected orally, inhibitory if they projected aborally. Circumferential was assigned based on the dominant orientation angle; these neurons typically arborized in myenteric ganglia, but may have traversed through the circular muscle. Epithelium-projecting neurons were not fully traced, given that peeled wholemount preparations were used. Neurons were assigned as epithelium-projecting if their fibres went into the circular muscle but did not arborize or clearly terminate there; typically only the first ∼100-150 µm of the neuron fibre was visible before it passed through the circular muscle.

#### Varicosity analysis

Neuronal varicosities were analyzed using the Spots module in Imaris 9.7. Varicosities were identified using syn-tdT expression, setting varicosity XY size to 1.2 µm diameter and modelling the point spread function elongation in the Z-axis as 4 µm diameter to avoid mistakenly stacking synapses. Spots were first filtered on Quality (based on syn-tdT fluorescence intensity); this identified all syn-tdT varicosities in the image, regardless of coexpression with other markers. Subsequent filtering identified only varicosities that colocalized with GFP fluorescence, excluding all others, to measure the total number of varicosities. This could then be further filtered based on fluorescence signal intensity of other markers, such as VGLUT2 IHC to determine whether varicosities were glutamatergic. Varicosity identification was manually checked to prevent the mis-assignment of GFP-negative varicosities to GFP-positive neurons, such as in the case of overlapping fibres. Varicosities were assigned to the primary fibre (the longest continuing fibre at a branch point) or branches manually, with the start and end of the primary axon fibre being defined as the first or last ∼200 µm of the fibre.

#### Output analysis

Recipient neurons were assigned if they were within 1 µm of a varicosity of minimum size 1 µm, and manually assigned using the Spots module in Imaris 9.7. If a recipient neuron did not express a marker, it was marked as HuC/D-only. Typically 20-30 branches were analyzed per neuron; the mean proportion of output neurons identified as a given type was calculated and presented per neuron, with a minimum of 4 neurons per mouse. Data was collated and analyzed in Microsoft Excel.

### Video analysis

Videos of optogenetic stimulation of *ex vivo* colons were analyzed in ImageJ/FIJI, with individual trials being treated independently. Videos were split into pre-stimulation and post-stimulation periods, ignoring the 20 s stimulation period in between. Pre-stimulation covered the 2 minutes before the stimulation start time. Post-stimulation analysis covered between 30 s and 2 minutes after the stimulation end time; the analysis period stopped if the pellet did not move or stopped moving (<1 mm in 30 s) to avoid interpreting spontaneous movement as stimulation-induced. Measurements of colon length and pellet start and end points were taken manually in FIJI, while the TrackMate plugin^48^ (v7.11) for FIJI was used to analyse pellet movement before and after stimulation. A spot diameter of 4-6 mm was used to identify the pellet using the LoG detector, while tracks were analysed with the Overlap Tracker. The desired track was isolated through filtering by quality, location, and distance travelled, with further adjustments made using TrackScheme as needed.

Spot and edge data were exported to produce individual pellet tracks in R, normalising tracks such that 0-100% represented the full length of the colon. Plotted post-stimulation tracks were overlaid with pre-stimulation tracks in Illustrator. Track data was exported from Trackmate to Microsoft Excel for measures of distance and speed. Distance was calculated using manual start and end points, normalised to percentages. Speed was calculated by dividing track displacement by track duration. For the analysis of speed and distance by pre-stimulation activity, three categories were defined based on if there had been significant movement (>5 mm travelled) 0-15 s, 15-30 s, or 30+ s before stimulation.

Spatiotemporal maps (STMs) were generated using Scribble 2.0 and Matlab (2012a) plugin Analyze 2.0^45^.

PenkCre-VGLUT2Flx videos *ex vivo* colonic motility was analysed using TrackMate as described above, with trials averaged per mouse.

### Statistics

All statistical tests and graphical representation of data were performed using Prism 9 software (GraphPad), other than plotting artificial pellet tracks, which was performed in R. Statistical comparisons were performed using one-way ANOVA to determine significant differences in a number of parameters, including between regions, types of neuron, and synaptic innervation, with Tukey’s or Sidak’s multiple comparisons test being employed to further investigate differences between individual groups, depending on the comparison being performed. Two-way ANOVAs with Sidak’s multiple comparisons test were used where two factors grouped the data, including sex and genotype, genotype and stimulus condition, or stimulus condition and stimulus location. The Kruskal-Wallis test with Dunn’s multiple comparisons test was used to investigate differences in recipient neuron identity, given the non-parametric nature of the data. The results of these tests are indicated on graphs as asterisks to indicate significance of at least p<0.05. Chi-squared analysis was used to determine if there was an enrichment for cells of a specific neuronal subtype receiving input from VGLUT2 neurons when compared to their overall representation in the MP, for example if a neuron subtype was 10% of all MP neurons, but received 20% of the contacts from VGLUT2 neurons.

## Acknowledgements

We thank members of the Kaltschmidt lab for experimental advice and discussions. We thank Dr. Vanda A. Lennon (Mayo Clinic) for the human HuC/D primary antibody, and Dr. Nirao Shah for the Prlr-IRES-Cre mouse. We thank Dr. Beatriz Robinson and Dr. Lucy Xu for their assistance with data analysis. This work was supported by an EMBO Fellowship ALTF 180-2019 (R.H.), Bertarelli Foundation Fellowship (J.L.B.), NIH T32-MH020016 (J.L.B.; K.R.), Stanford Neurosciences Interdepartmental Graduate Program (K.R.), a research grant from The Shurl and Kay Curci Foundation (J.A.K.), the Firmenich Foundation (J.A.K.), the Carol and Eugene Ludwig Family Foundation (J.A.K.), Stanford ADRC Developmental Project Grant (National Institutes of Health Grant P30AG066515) (J.A.K.), National Institutes of Health Grant R21 HD110950 (J.A.K.), the Wu Tsai Neurosciences Institute (J.A.K.) and the Stanford University Department of Neurosurgery (J.A.K.).

## Author Contributions

R.H. and J.A.K designed and conceptualised the project. R.H., J.B. and K.R. performed the experiments. R.H. and J.B. analysed the experimental results. E.T.Z. provided resources. R.H. wrote the manuscript with feedback from all authors. J.A.K. supervised the project.

---

**Extended Data Fig. 1.**
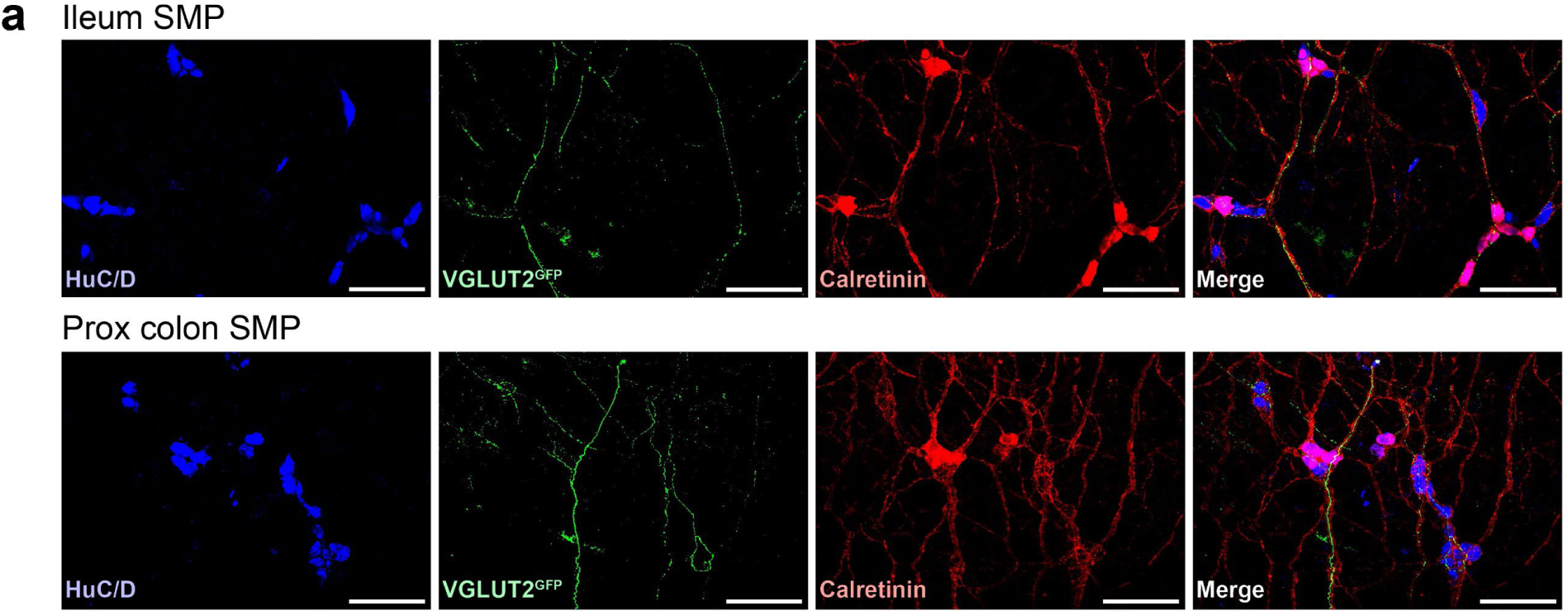
VGLUT2 expression in the submucosal plexus (SMP). **a**, Representative images of adult wholemount MP showing VGLUT2^GFP^ (green), calretinin (red), and neuronal label HuC/D (blue) in the SMP of the ileum (top) and proximal colon (bottom).

**Extended Data Fig. 2.**
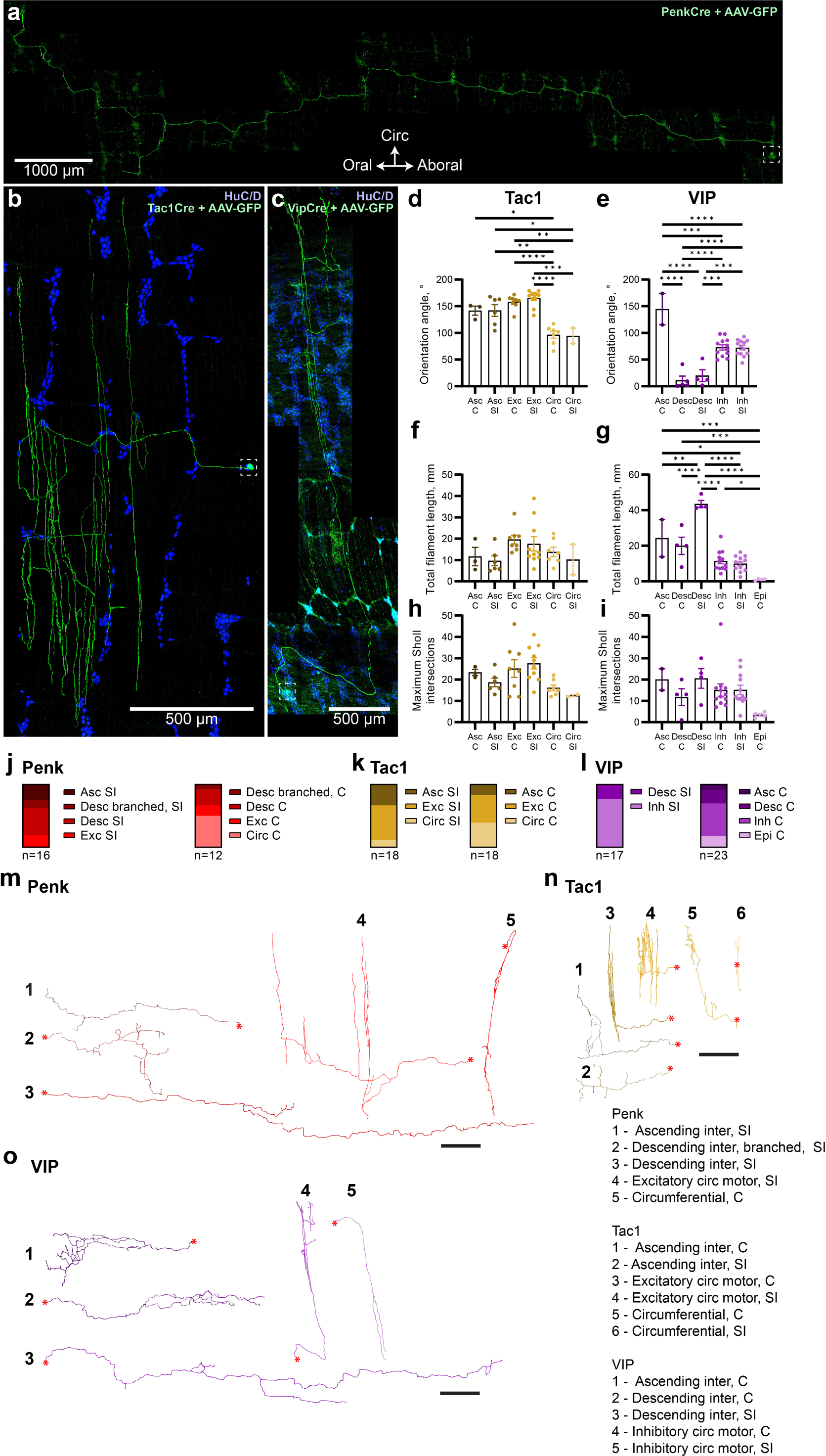
Neuronal morphology of other enteric markers reveals diversity of neuronal classes. **a-c**, Representative images of a Penk^AAV-GFP^ ascending interneuron (**a**), Tac1^AAV-GFP^ excitatory motor neuron (**b**) and Vip^AAV-GFP^ inhibitory motor neuron (**c**), which were transduced by Cre-dependent AAV-GFP and immunostained for GFP (green) and HuC/D (blue; **b,c** only). Neuronal somata indicated by white boxes. Scale bars as indicated. **d-i**, Quantification (mean ± SEM) of Tac1^AAV-GFP^ (**d,f,h**) and Vip^AAV-GFP^ (**e,g,i**) neuron orientation (**d,e**), total filament length (**f,g**), and maximum Sholl intersections (**h,i**). n=2-12 neurons per group. Each dot represents a different neuron, taken from across 3 (Tac1) and 7 (Vip) mice. At least two neurons had to be identified as a given classification to be included in this analysis. Penk^AAV-GFP^ neurons were not analysed in this way due to the large diversity of different neuron classifications and resulting low n per class. All tests one-way ANOVA. *p< 0.05, **p< 0.01, ***p< 0.001, ****p< 0.0001. **j-l**, Proportions of different neuron classes identified for Penk^AAV-GFP^ (**j**), Tac1^AAV-GFP^ (**k**), and Vip^AAV-GFP^ (**l**) in the SI (left) and colon (right). n as indicated. At least two neurons had to be identified as a given classification to be included in this analysis. **m-o**, Representative traces of Penk^AAV-GFP^ (**m**), Tac1^AAV-GFP^ (**n**), and Vip^AAV-GFP^ (**o**) neuron classes. Numbers beside each trace correspond to legend (bottom right). Scale bars: 1000 µm for motor and circumferential neurons; 2000 µm for interneurons. Soma location indicated by red asterisk. Abbreviations for **d-l**: C: colon; SI: small intestine; Asc: ascending interneuron; Desc: descending interneuron; Exc: excitatory motor neuron, Circ: circumferential neuron; Inh: inhibitory motor neuron; Epi: epithelium-projecting neuron. Note that for epithelium-projecting neurons, only the portion within the muscularis was traced and measured.

**Extended Data Fig. 3.**
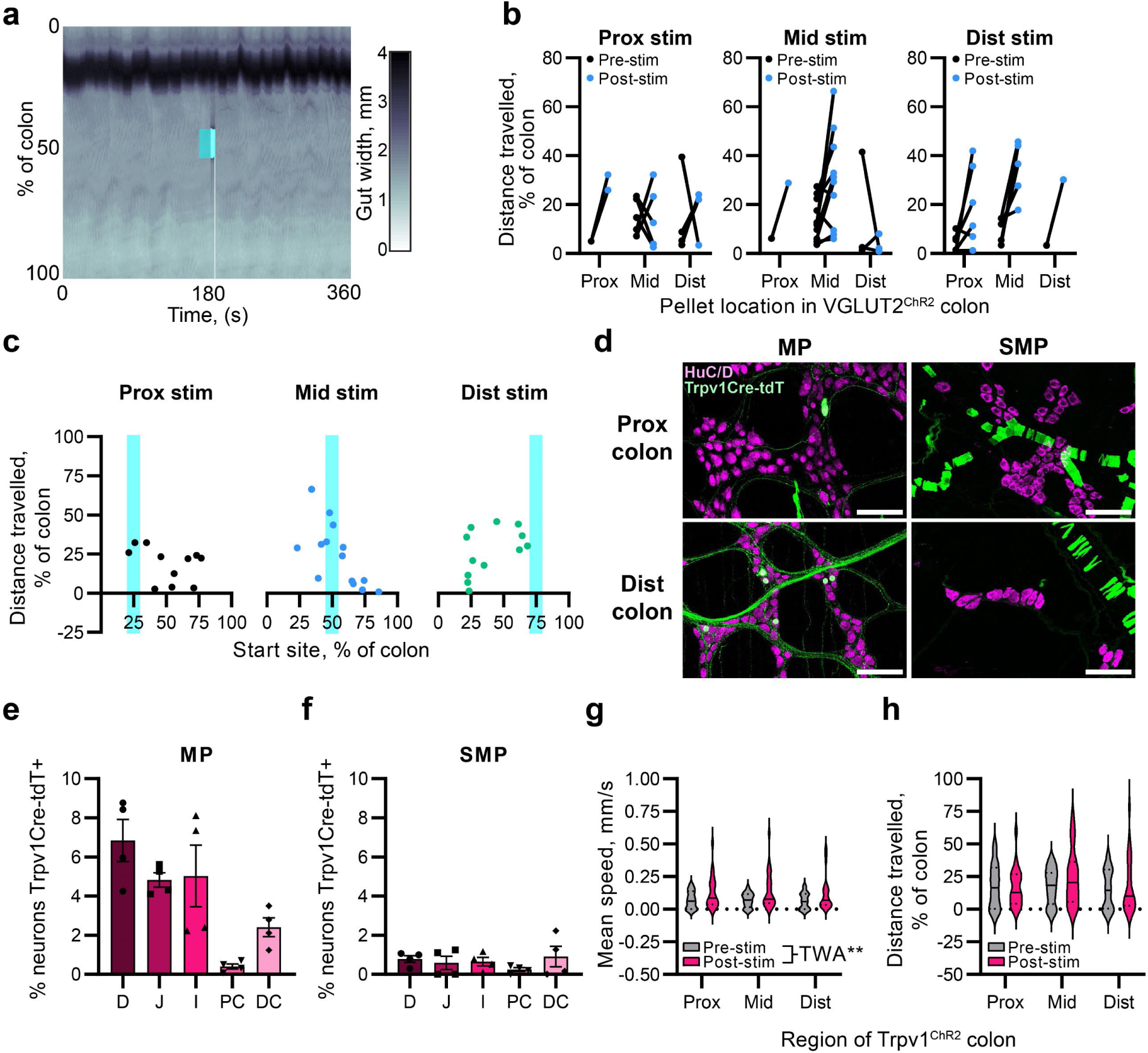
Optogenetic excitation of VGLUT2, Trpv1 and Vip populations. **a**, Representative spatio-temporal map of a VGLUT2^ChR2^ colon treated with 1 µm tetrodotoxin (TTX), showing the width of each point along the length of the colon (y axis, %) over 6 minutes. Optogenetic stimulation (cyan box) occurs half way through the recording. The dark band indicates the artificial faecal pellet. **b**, Distance travelled by artificial pellets in 2 minutes before optogenetic stimulation, and up to 2 minutes after stimulation of VGLUT2^ChR2^ colons, split by pellet location and stimulation location. **c**, Correlation between distance travelled by artificial pellets as a proportion of the total length of colon and the location of the pellet at the time of stimulation (start site), split based on stimulation location (cyan). Each dot represents a single trial. Graphs are alternate visualisations of Fig. 5j. **d**, Representative images of adult wholemount MP showing Trpv1^tdT^ (green) and neuronal label HuC/D (magenta) in the proximal colon (top) and distal colon (bottom) MP (left) and SMP (right). Scale bars 100 µm. **e,f**, Proportion of total HuC/D neurons (mean ± SEM) positive for Trpv1^tdT^ in the MP (**e**) and SMP (**f**) across intestinal regions. **g,h**, Violin plots of artificial pellet speed (**g**) and distance travelled (**h**) following mid-colon optogenetic stimulation of Trpv1^ChR2^ colons, divided by stimulation location. Two-way ANOVAs (TWA) performed for pre/post stimulation and stimulation location. Comparisons in which there was an overall significant effect of stimulation but no significant differences following multiple comparisons testing are indicated (**i**). *p< 0.05, **p< 0.01, ****p< 0.0001. Abbreviations: D: duodenum; J: jejunum; I: ileum; PC: proximal colon; DC: distal colon.

**Extended Data Fig. 4.**
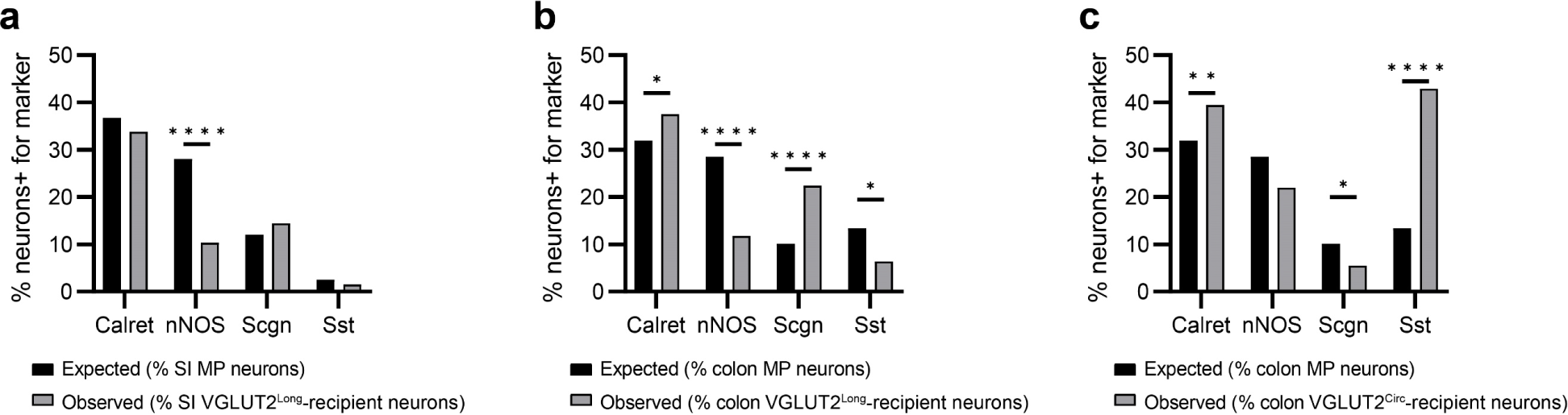
Chi-squared analysis of VGLUT2 neuron recipient populations. **a-c**, Proportion of total small intestine (**a**) or colon (**b,c**) myenteric neurons expressing a given marker (calretinin, nNOS, Scgn or Sst) compared with the proportion of neurons receiving input from VGLUT2^Long^ (**a,b**) or VGLUT2^Circ^ (**c**) neurons that express the marker. All analyses chi-squared test, *p< 0.05, **p< 0.01, ****p< 0.0001.

**Extended Data Fig. 5.**
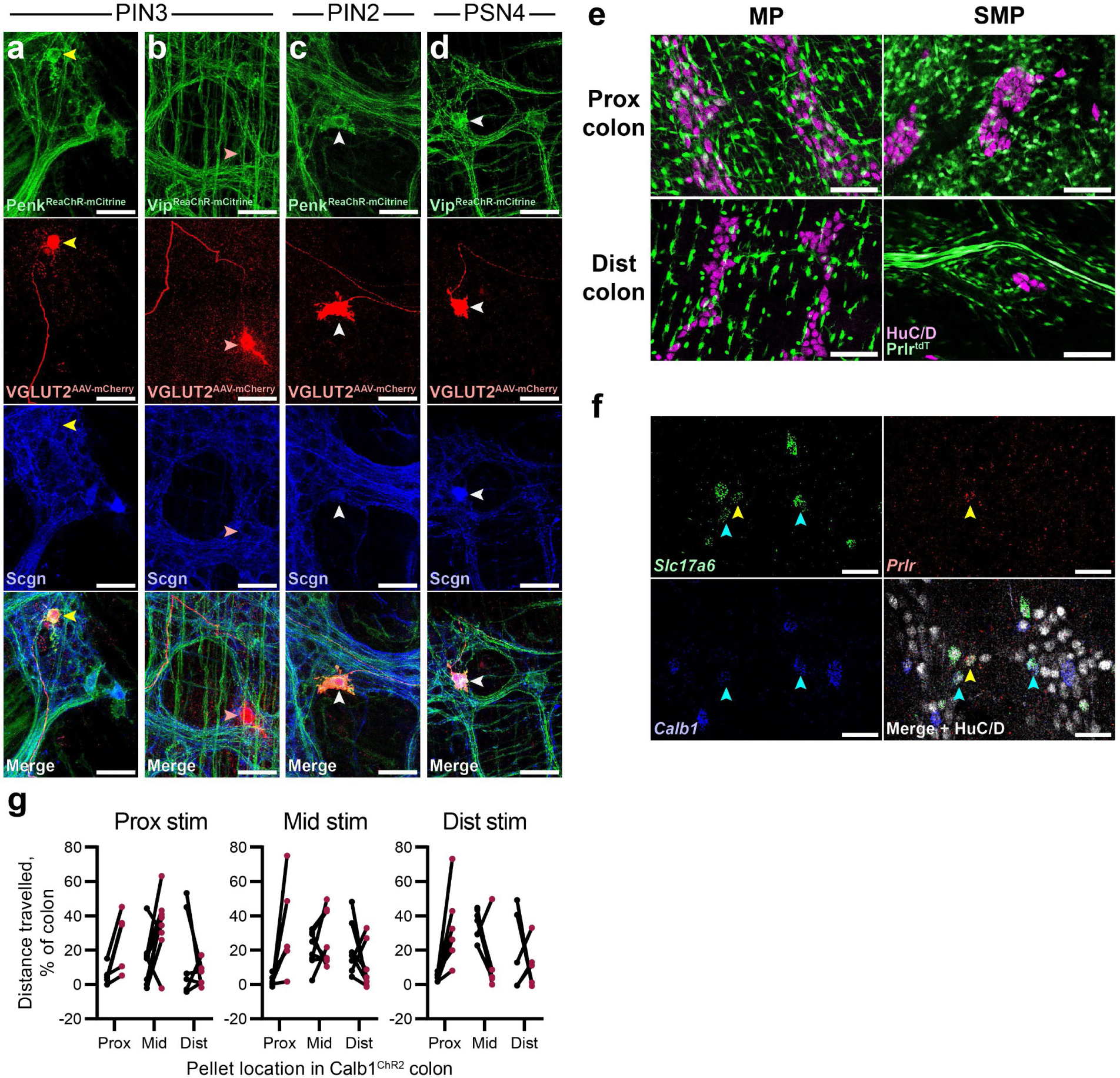
Prlr and Calb1 as candidate VGLUT2 neuron population markers. **a-d**, Representative images of adult wholemount MP showing VGLUT2^AAV-mCherry^/Penk^ReaChR-mCitrine^ (**a,c**) and VGLUT2^AAV-mCherry^/Vip^ReaChR-mCitrine^ (**b,d**) to determine colocalisation between sparsely transduced VGLUT2 neurons (red), mCitrine (green) and Scgn IHC (blue) for the assignment of VGLUT2^Circ^ and VGLUT2^Long^ neurons to one of three previously identified ENS clusters: putative interneuron 2 (PIN2), PIN3, and putative sensory neuron 4 (PSN4). Yellow arrowheads show VGLUT2^AAV-mCherry^ colocalisation with mCitrine without Scgn; red arrowheads show VGLUT2^AAV-mCherry^only; white arrowheads show colocalisation between VGLUT2^AAV-mCherry^, mCitrine and Scgn. **e**, Representative images of adult wholemount MP showing Prlr^tdT^ (green) and neuronal label HuC/D (magenta) in the proximal colon (top) and DC (bottom) MP (left) and SMP (right). **f,** Representative images of adult wholemount MP showing Slc17a6 (VGLUT2; green), Prlr (red) and Calb1 (blue) RNAscope and alongside HuC/D (grey) IHC in the proximal colon. Cyan arrowheads indicate *Calb1*+/*Slc17a6*+ neurons, yellow arrowheads indicate *Prlr*+/*Slc17a6*+ neurons. **g**, Distance travelled by artificial pellets in 2 minutes before optogenetic stimulation, and up to 2 minutes after stimulation of Calb1^ChR2^ colons, split by pellet location and stimulation location. Scale bars: 50 µm (**a-d,f**), 100 µm (**e**).

